# Natural Language Processing-like Deep Learning Aided in Identification and Validation of Thiosulfinate Tolerance Clusters in Diverse Bacteria

**DOI:** 10.1101/2024.09.03.611110

**Authors:** Brendon K. Myers, Anuj Lamichhane, Brian H. Kvitko, Bhabesh Dutta

**Author notes:** Corresponding Authors: Bhabesh Dutta, Brian H. Kvitko, **Email:**. **Author Contributions:** BKM conducted all computational analysis. AL designed the gene validation experiments. BHK offered expertise in allicin tolerance, bacterial genetics, and protein biochemistry. BD funded the project. BKM and BD designed the project. BKM wrote the initial drafts of the manuscript. AL, BHK, and BD provided input on the manuscript. **Competing Interest Statement:** The authors declare that they have no known competing financial interests or personal relationships that could have appeared to influence the work reported in this paper.

## Abstract

Allicin tolerance (*alt*) clusters in phytopathogenic bacteria, which provide resistance to thiosulfinates like allicin, are challenging to find using conventional approaches due to their varied architecture and the paradox of being vertically maintained within genera despite likely being horizontally transferred. This results in significant sequential diversity that further complicates their identification. Natural language processing (NLP) - like techniques, such as those used in DeepBGC, offers a promising solution by treating gene clusters like a language, allowing for identifying and collecting gene clusters based on patterns and relationships within the sequences. We curated and validated *alt*-like clusters in *Pantoea ananatis* 97-1R (PA), *Burkholderia gladioli* pv. *gladioli* FDAARGOS 389 (BG), and *Pseudomonas syringae* pv. tomato DC3000 –(PTO). Leveraging sequences from the RefSeq bacterial database, we conducted comparative analyses of gene synteny, gene/protein sequences, protein structures, and predicted protein interactions. This approach enabled the discovery of several novel *alt*-like clusters previously undetectable by other methods, which were further validated experimentally. Our work highlights the effectiveness of NLP-like techniques for identifying underrepresented gene clusters and expands our understanding of the diversity and utility of *alt*-like clusters in diverse bacterial genera. This work demonstrates the potential of these techniques to simplify the identification process and enhance the applicability of biological data in real-world scenarios.

**Significance Statement:** Thiosulfinates, like allicin, are potent antifeedants and antimicrobials produced by *Allium* species and pose a challenge for phytopathogenic bacteria. Phytopathogenic bacteria have been shown to utilize an allicin tolerance (*alt*) gene cluster to circumvent this host response, leading to economically significant yield losses. Due to the complexity of mining these clusters, we applied techniques akin to natural language processing to analyze Pfam domains and gene proximity. This approach led to the identification of novel *alt*-like gene clusters, showcasing the potential of artificial intelligence to reveal elusive and underrepresented genetic clusters and enhance our understanding of their diversity and role across various bacterial genera.

## Introduction

Plants deploy an impressive array of small molecules to defend themselves against herbivory and pathogen-mediated infection. Thiosulfinates such as allicin are charismatic small molecules known for their role as antifeedants and antimicrobials in *Allium* species (1, 2). These small molecules are reactive organosulfur compounds responsible for several *Allium* species’ characteristically pungent flavor and smell (3, 4). Allicin is produced when the enzyme alliinase acts on alliin, transforming it into a thiol-reactive compound. It interacts with cellular thiols, leading to allyl-mercapto modifications in proteins that deactivate enzymes and cause protein aggregation (5). Additionally, allicin reacts with reduced glutathione, converting it to S-allylmercaptoglutathione and thus depleting the cellular glutathione pool (5, 6). Thiosulfinates have been demonstrated to be inhibitory to a wide range of microorganisms both *in vitro* and *in vivo* (7, 8, 9, 10). Recently, gene clusters associated with allicin tolerance were identified not only in the onion pathogens *Pantoea ananatis* (PA) and *Burkholderia gladioli* (BG) but also in the garlic saprophyte *Pseudomonas fluorescens* (10, 11, 12). These genes were named allicin tolerance (*alt*) genes and are enriched for genes involved in thiol redox reactions. The *alt* clusters increased onion virulence capacity in PA and BG strains and conferred increased allicin tolerance to *E. coli* (12, 13). The *alt* gene cohort appears to function additively for managing cellular thiol stresses, with multiple genes conferring partial tolerance phenotypes (10). In their 2018 study, Stice et al. (10) data mined the NCBI GenBank database to identify *Pantoea* spp. with *alt* clusters, using the *altG* gene as an indicator. In doing so, the authors observed that several strains isolated from *Allium* hosts and some *Brassica* species carry *alt* clusters. In contrast, strains isolated from non-thiosulfinate-producing hosts did not. Inspired by an intuitive understanding of the characteristics defining an *alt* cluster, manual curation led to discovering a unique cluster within BG and other *Burkholderia* spp. (12), supported by multigene BlastX analysis (12).

Considering the importance of *alt* clusters in thiosulfinate tolerance and plant-microbe interactions, identifying the variety of *alt* clusters and their presence in bacterial species is crucial. The *alt* clusters that were characterized and validated share little sequence or gene synteny similarity. Typical gene-mining techniques, such as NCBI BLAST or multigene BLAST, do not translate well between *alt* clusters localized within different bacterial genera. Although the *alt* cluster is potentially horizontally transferred as it is localized on plasmids, it seems to be maintained vertically within individual bacterial genera. This makes identifying *alt* clusters within genera comparatively easier; however, their identification among distinct genera is quite challenging. Isolating thiosulfinate-tolerant bacteria from a thiosulfinate-producing host and then manually curating the annotations list for a conspicuous gene cluster has been the *modus operandi* for *alt* gene cluster discovery to date. However, it is a time-consuming process that requires in-depth training and a reliable annotation pipeline. Even in optimal conditions, individual researchers might develop personal biases towards which annotations they deem more reliable or questionable, potentially resulting in misidentification of *alt* clusters. To formalize an *alt*-identification and recovery method independent of the issues caused by gene sequence and gene synteny, we used NLP-like techniques for mining putative *alt*-like gene clusters.

The methodology employed here is similar to those used with genome mining for secondary metabolite biosynthetic gene clusters. These pipelines must overcome a challenging task that requires careful consideration of gene content. For example, bacteria tend to organize genes into localized clusters to make metabolite synthesis more efficient (14,15,16). While manual curation and BLAST are effective for similar biosynthetic gene clusters (BGCs) in closely related organisms, they fall short when sequence data alone is inadequate or manual efforts are impractical due to time or cost (17). In such cases, more rigid, ‘hard-coded’ algorithms are used, though they require predefined gene and protein data rules, limiting their use with less-defined gene clusters (18, 19). Machine learning is the natural next step in algorithmic complexity to solve these problems, autonomously allowing for a more generalizable “learning” of input content. This allows for discovering more novel BGC’s as the algorithm generates its own rules during training for further downstream applications.

An example of this is ClusterFinder (20). ClusterFinder utilizes a Hidden Markov Model (HMM) approach rather than sequence alignment, allowing for greater freedom of discovery. However, HMM does not preserve position dependency effects or any potential higher-order information that may be relevant for BGC discovery (21, 22, 23).

To address the need for higher-order information in BGC discovery, a deep learning approach using Recurrent Neural Networks (RNNs) with the addition of vector representations of protein family tags (Pfam) was designed, which improved the capacity for algorithmically detecting novel BGCs (24). DeepBGC utilizes an NLP strategy for identifying and even extracting novel BGCs from bacterial genomes via a clever use of a Bidirectional Long Short-Term Memory (BiLSTM) RNN (25,26) and a word2vec-like word embedding skip-gram neural network that the authors named pfam2vec (24).

In this work, we trained DeepBGC on our small collection of validated *alt* clusters to determine the potential for more complex artificial intelligence methods to accelerate the discovery process. Although the *alt* cluster does not represent a typical BGC where each gene collaboratively synthesizes a molecule, the organization and perceived additive function of these genes for the *alt* phenotype renders the cluster amenable to methodologies like those used in BGC discovery. The new *alt* model was then utilized to data mine the entire RefSeq bacterial database for potential *alt*-like clusters. Representative clusters were selected and refined through manual curation and sequencing data analysis to produce representative sequences of *alt*-like gene clusters. The genes, proteins, and predicted binding potential for selected genes from each cluster were compared to identify potentially valuable methodologies for differentiating *alt*-like gene clusters. Finally, chosen *alt*-like gene clusters were validated by expression of synthesized gene pairs in PA PNA 97-1 Δ*alt* and screening for increased thiosulfinate tolerance based on the improved ability of strains to grow in thiosulfate-rich onion extract.

## Results

### *alt* Seed Cluster Gene and Protein Sequence Comparisons Show Low Sequence Similarity

Onion-associated bacteria like PA and BG possess *alt*-clusters that impart the ability to survive and propagate in thiosulfinate-rich-environments (figure 1A). In some cases, this may lead to bulb rot symptoms (figure 1A). In the genomic comparison of PA, PTO, and BG, the total gene counts are 11, 16, and 7, respectively. Shared genes across these strains include *altA*, *altB*, *altC*, *altE*, *altR*, *altJ*, and *altI*. When evaluating synteny, among PA, PTO, and BG there appears to be little in common between the three sequences. Between PA and BG, *altA* and *altC* do localize; however, their order is inverted. Further, the *altR* and *altE* are adjacent between both PA and BG. The *altJ* and *altB* are adjacent but inverted between BG and PTO. The *altE* and *altA* are also adjacent but inverted between BG and PTO (figure 1B).

**Figure 1.**
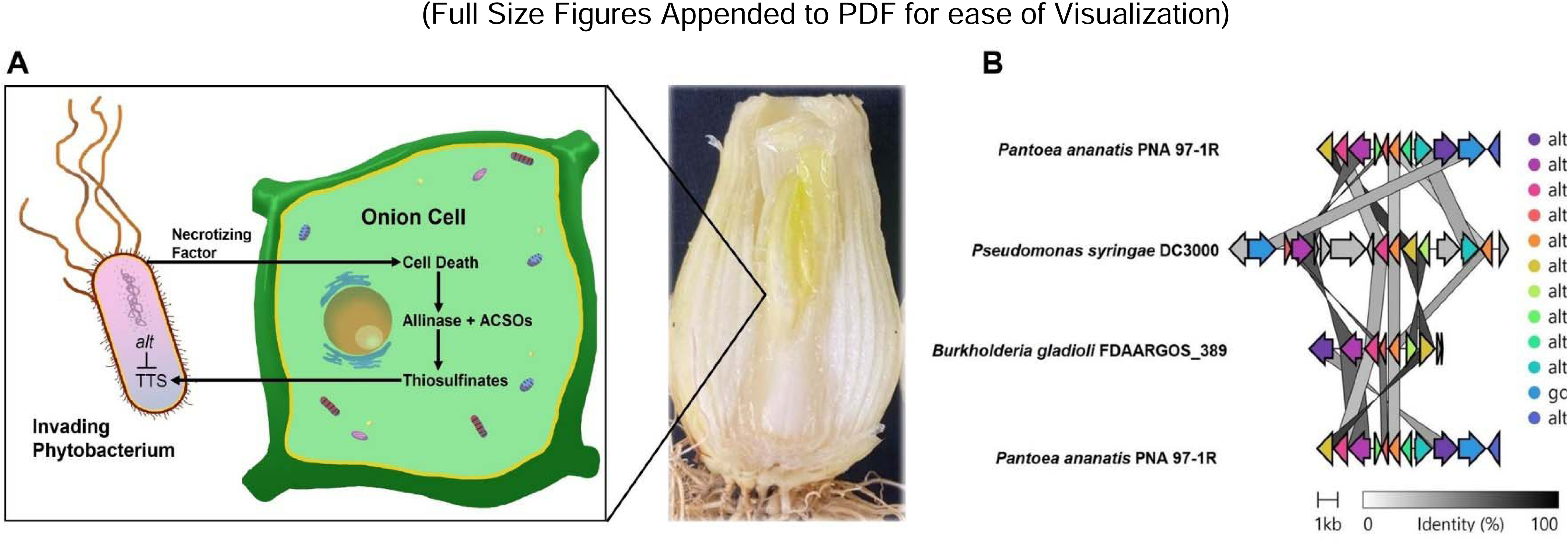
Overview of the importance of thiosulfinate tolerance with *Pantoea ananatis* as a model example. (A) Pictorial representation of the chemical arms race between an invading phytobacterium and its *Allium* host, depicted as *Allium cepa*. When the phytopathogen utilizes its necrotizing factors to kill the host cells, it, in turn, becomes challenged with toxic thiosulfinate stress (TTS) that is managed by the allicin tolerance (*alt*) cohort. In addition, an example of bulb-rot symptoms due to *Pantoea ananatis* compatible interactions in *A. cepa* in onion bulbs is included. The provided example is a longitudinal section of an infected bulb displaying rotten water-soaked center scales with visible bacterial growth. (B) Gene cluster synteny comparisons of *alt* clusters used as the input sequences for DeepBGC training. These comparisons were generated with Clinker. The arrows represent coding sequences along with their directionality. Shaded lines reflect the degree of similarity between the gene clusters, with darker shades indicating higher similarity. Arrows are colored based on their *alt* annotations.

We observed high degrees of dissimilarity when comparing the total gene cluster sequence similarity among our original three validated *alt* clusters. Additionally, when analyzing individual genes with annotations shared across all three clusters, the similarity percentages exhibit a range between 21.9 and 74.1%. Specifically, *altI* sequences show similarities from 39.1 to 52.1%, *altA* from 66 to 69.9%, and *altC* from 47.3 to 50.6%. Sequences of *altE* vary from 62 to 69.4%, *altR* from 46.5 to 51.2%, and *altJ* from 41.1 to 70.5%. The *altB* gene maintains high consistency around 74% across all comparisons. A second *altR* gene in the PTO cluster displays 47.5 to 54.5% similarity. For genes only shared between PA and PTO, the lowest similarity is noted in *altJ* at 21.9%, with other genes like *altD*, *altH*, and gor displaying up to 52.4% similarity (figure 1, table 1).

**Table 1.**
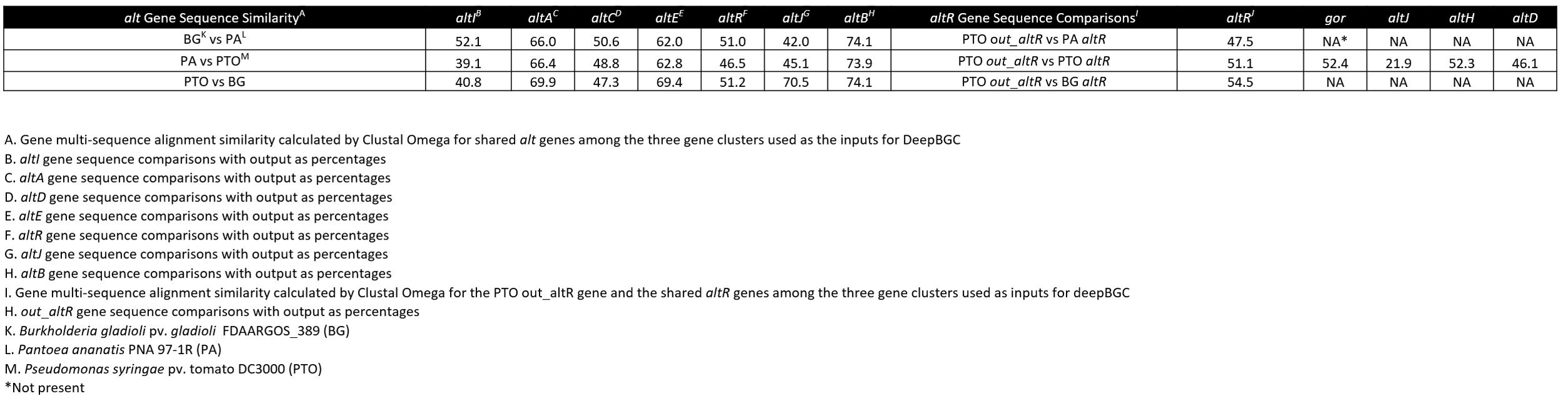
Multi-sequence alignment (MSA) comparison of allicin tolerance (*alt*) genes among shared genes in the *alt* clusters used as the inputs for training by DeepBGC. Sequences were retrieved from *Burkholderia gladioli* pv. *gladioli* FDAARGOS_389 (BG), *Pantoea ananatis* PNA 97-1R (PA), and *Pseudomonas syringae* pv. tomato DC3000 (PTO). MSA was calculated using Clustal Omega in the default settings at https://www.ebi.ac.uk/Tools/msa/clustalo/. Comparisons were made between shared alt genes (*altI*, *altA*, *altC*, *altE*, *altR*, *altJ*, and *altB*) as well as an additional comparison between a secondary *altR*-like gene in *P. syringae* pv. tomato DC3000 (*out*_*altR* as in figure 1) between the three bacterial strains that were used as input data in DeepBGC.

A broad range of dissimilarities is observed in assessing protein sequence similarity across the three validated alt clusters, with percentages ranging from 18.1% to 82.1%. Notably, *altI* shows significant variation, with 48.2% similarity between PA and BG, dropping to 18.1% when comparing BG vs. PTO. The protein sequences in *altA* range from 67.9 to 74.3% across comparisons, while *altC* varies from 38.5 to 43.8%. The *altE* sequences are relatively similar, ranging from 62.5 to 71.8%. The *altR* protein sequences vary from 35.4 to 43.9%, and *altJ* from 27.2 to 76.6%. The *altB* exhibits high consistency, with similarities ranging from 78.5 to 82.1%. A second *altR* in the PTO cluster shows similarities between 36.5% to 47.3%. Similarities for proteins exclusively shared between PA and PTO are notably lower, with *altJ* at 25.5%, *altD* at 30.5%, *altH* at 45.5%, and *gor* at 43.9% (table 2).

**Table 2.**
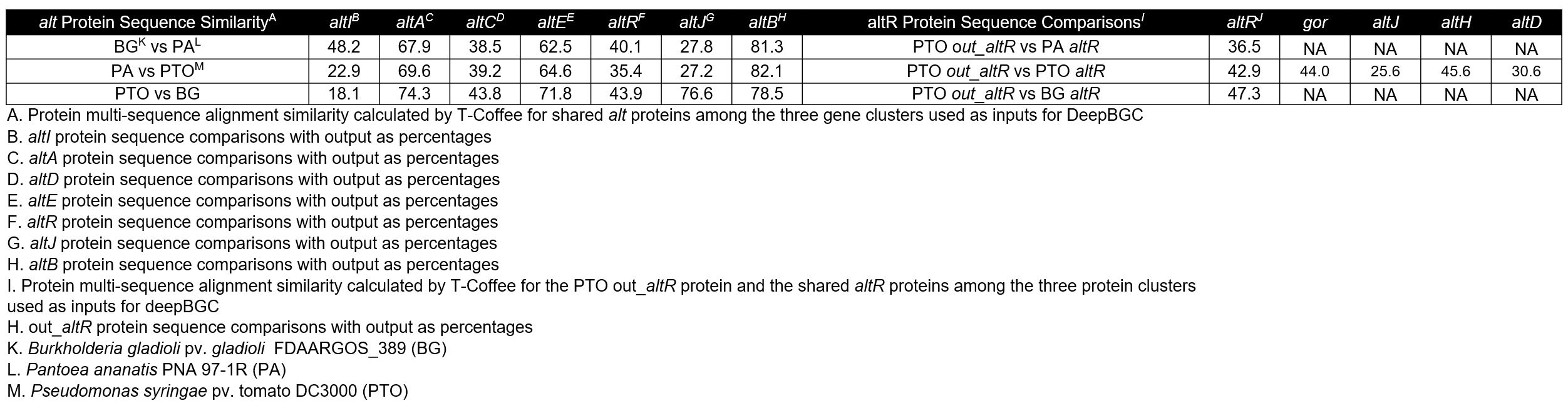
Multi-sequence alignment (MSA) comparison of allicin tolerance (*alt*) proteins among shared proteins in the *alt* clusters used as the inputs for training by DeepBGC. Sequences were retrieved from *Burkholderia gladioli* pv. *gladioli* FDAARGOS_389 (BG), *Pantoea ananatis* PNA 97-1R (PA), and *Pseudomonas syringae* pv. tomato DC3000 (PTO). MSA was calculated using T-Coffee in the default settings at https://www.ebi.ac.uk/Tools/msa/tcoffee/. Comparisons were made between shared *alt* proteins (*altI*, *altA*, *altC*, *altE*, *altR*, *altJ*, and *altB*) as well as an additional comparison between a secondary *altR*-like protein in *P. syringae* pv. tomato DC3000 (out_*altR* as in figure 1) between the three bacterial strains that were used as input data in DeepBGC.

### DeepBGC Data Mining of the NCBI RefSeq Database and Filtering for Autonomous Collection of *alt*-like Gene Clusters

DeepBGC training on the three validated *alt* cluster sequences was repeated 15 times, and the reports were compared to assess model performance. The average loss across runs was minimal at 0.00, indicating steady performance. However, a maximum loss value of 0.40 suggests some performance variability. In assessing model performance for DeepBGC, accuracy was consistent across all tests, averaging 1.00 with a standard deviation of 0.00 and a minimum accuracy of 0.98, highlighting the model’s reliability. Precision and recall were both low, averaging 0.01, indicating a challenge in accurately identifying and capturing true positives from the dataset. This variability was reflected in the AUC-ROC scores, which averaged 0.82, suggesting good discriminatory ability with room for improvement. Statistical analysis confirmed significant variability in precision (F-value: 3.78, p-value: ∼2×10^−6^), recall (F-value: 5.17, p-value: ∼7.7×10^−10^), and AUC-ROC (F-value: 16.09, p-value: ∼7.64×10^−43^). In contrast, differences in loss and accuracy were not statistically significant (F-values: 0.86 and 0.39, p-values: 0.60 and 0.98, respectively), indicating stable performance in these areas. The detailed statistical insights underscore the need for further refinement to enhance precision, recall, and overall model robustness. These results are expected with the small training dataset we can access and are overcome with manual inspection of DeepBGC extractions for validity.

Upon completion of the DeepBGC-enabled data mining of 238,362 bacterial genomes from RefSeq, we extracted 12,280 gene clusters. These were reduced to 1,777 sequences post MMseqs2 redundancy filtering with an average GC% of 53.5% (median 55.1%, max. 76.6% and min. 25.7%), an average sequence length of 8,800 (max 61,726, min 1,215, and median of 6,424), and finally an average file size of 23KB (max 114KB, min 9KB, and a median of 19KB). After further manual curation to remove all gene sequences that appear split by the end of contigs, only four genes in total length, or do not have at least 3 unique *alt*-like pfam tags, we chose 47 representative *alt*-like sequences. These 47 representative clusters contained an average GC% of 51.7% (max 69.9%, min 32.4%, and median 53.5%), an average sequence length of 7,931 (max 30,170, min 3,109, and median 6,316), and finally an average file size of 28 KB (max 114 KB, min 12 KB, and median 23 KB). When screening for clusters that are representative of our initial three *alt* clusters, the *Pantoea alt* cluster is represented by an *alt*-like cluster from *Duffyella gerundensis* (NZ_LN907829.1) with 94% sequence identity and identical values of assigned Pfam domains, the *Burkholderia alt* cluster is represented by a truncated *alt*-like gene cluster from *Paraburkholderia graminis* (NZ_CP024936.1) with 74% sequence similarity. The *Pseudomonas alt* cluster is represented by itself as PTO (NC_004578.1). For all downstream gene cluster comparisons, we used the PA and BG *alt* clusters for comparison as references (figure 2).

**Figure 2.**
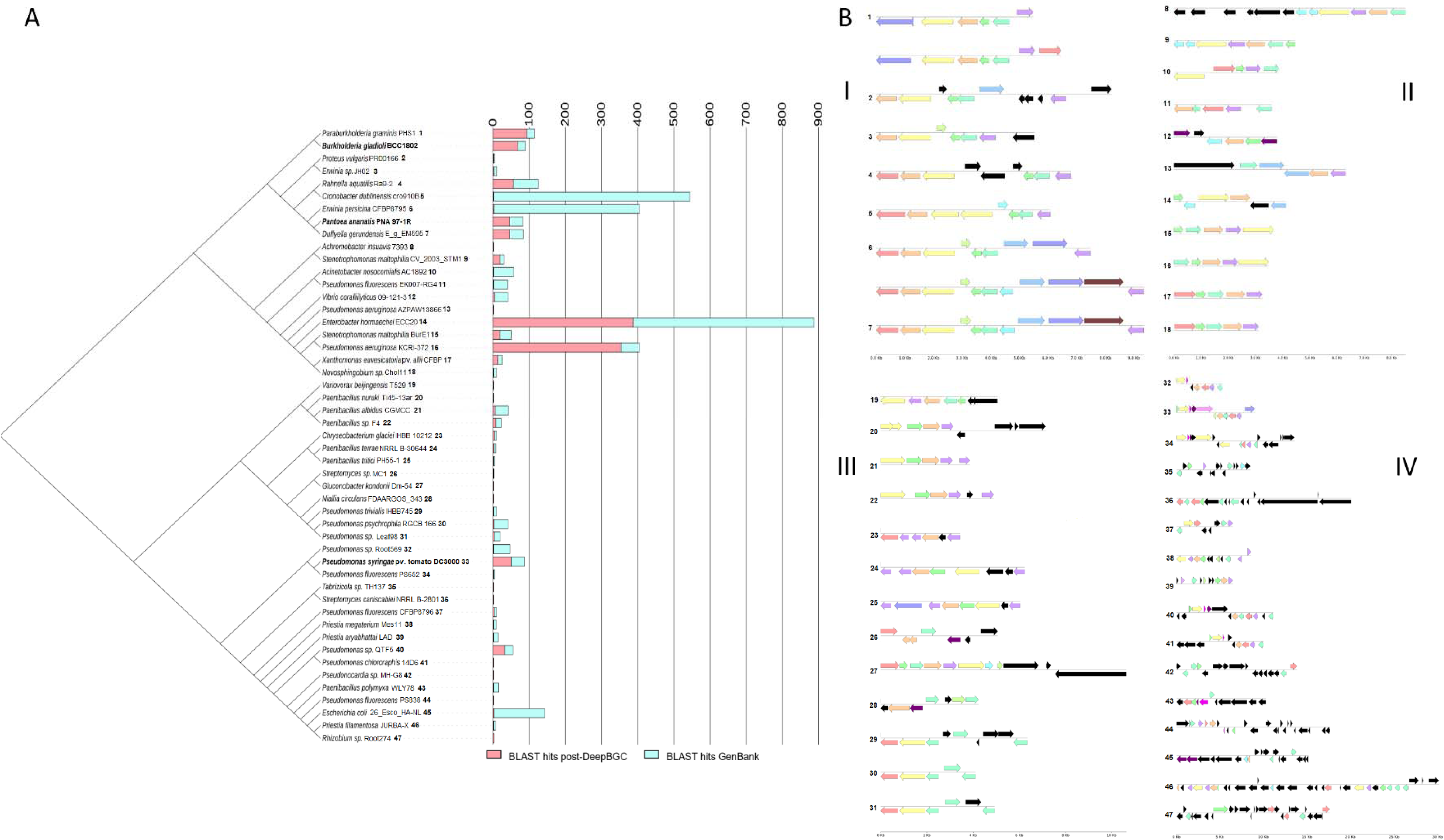
Allicin tolerance (*alt*)-like representative clusters post-DeepBGC extraction of the RefSeq bacterial database. (A) A comprehensive insight into the distribution of *alt*-like clusters within the NCBI system using the Levenshtein distance matrix of color-coded Pfam domain tags. The resulting BLAST hits of representative *alt*-like clusters on all extracted *alt*-like gene clusters from the RefSeq database are shown in pink, while the resulting BLAST hits of representative *alt*-like clusters from the online GenBank bacterial database are shown in blue. Each line indicates 50 sequences. This tree compares the pattern of Pfam tags in gene clusters and should not be misconstrued as a phylogenetic tree. (B) Color-coded examples of selected representative *alt*-like Pfam domain tags in the deepBGC-extracted clusters; (I) represents the first terminal group between *Paraburkholderia graminis* (PHS1) to *Duffyella gerundensis* (E_g_EM595); (II) represents the terminal group between *Achromobacter insuavis* (7393) to *Novosphingobium sp.* (Chol11); (III) represents the terminal group between *Variovorax bejingensis* (T529) to *Pseudomonas sp.* (Leaf98); (IV) represents the terminal group between *Pseudomonas spp.* (Root569) to *Rhizobium sp.* (Root274). These are unrooted neighbor-joining trees based on the Levenshtein differences between a color code conversion of Pfam tags into text strings using the Levenshtein python package. Gene clusters are numbered for ease of comparison. Gene clusters with similar Pfam annotation synteny, but different sequence content was listed as separate clusters (see clusters 17, 18).

Gene sequence similarity among these 49 clusters is low for alignment-based comparison methods. To minimize this, we color-coded genes based on their known relevance and converted the color code into strings for Levenshtein comparisons. These gene clusters are separated into 4 distinct groups (figure 2). The total counts for *alt*-like genes among these representative clusters are as follows, *altR* (N=41), *altC* (N=38), *altJ* (N=36), *altE* (N=36), *altA* (N=33), *altB* (N=28), *altG* (N=11), *altI* (N=8), *altD* (N=8), *PSPTO_4258* (N=7), *altH* (N=6), *PSPTO_4257* (N =6), *gor* (N =2), *kefC* (N=2), *PSPTO_5268* (N=1). Among these, *altR* has the highest count per gene cluster (figures 2 and 3).

### BLAST of DeepBGC-Mined *alt*-like Clusters and NCBI GenBank for Representative Sequence-Species Diversity Shows Wide Diversity of *alt*-like Gene Clusters Among Bacterial Genera

To compare the diversity of bacterial species represented by recovered *alt*-like gene clusters, we employed BLAST to retrieve clusters from both the sequences obtained through DeepBGC-enabled data mining of RefSeq and NCBI GenBank. Due to the varying selection of available sequences between NCBI’s RefSeq and GenBank, cross-comparison between the two databases may offer a more comprehensive understanding of species diversity compared to solely re-screening NCBI RefSeq with BLAST. Notably, *Klebsiella pneumoniae* emerged as a predominant species, constituting 56% of the recovered sequences in one instance and demonstrating significant representation across multiple samples. Conversely, specific sequences lacked a single dominant species, particularly those associated with *Stenotrophomonas maltophilia*. Detailed analysis of biodiversity using Shannon-Wiener indices unveiled varying levels of diversity among samples. For instance, sequences attributed to *S. maltophilia* exhibited higher diversity, representing 63-84% of recovered sequences. In contrast, sequences linked to *Pseudomonas aeruginosa* displayed lower diversity, comprising 77-82% of sequences. Additionally, GenBank BLAST analysis yielded taxonomic insights into the retrieved sequences. While *Klebsiella pneumoniae* was prevalent, other species, such as *Pseudomonas fluorescens* and *Escherichia coli*, were also prominently featured. Specific genera exhibited species-specific enrichment, with *Pseudomonas* and *Paenibacillus* showing pronounced representation in the sequences. These findings underscore the wide distribution of *alt*-like gene clusters across bacterial species and highlight their potential ecological importance (figure 2). These results are summarized in the supplementary table (supplementary table 1).

### 3D Superimposition of Predicted Protein Models is Insufficient for Differentiating Between alt, alt-like, and Unrelated Proteins

Due to the complexity inherent in classifying *alt* clusters by sequence and gene synteny, we investigated potential discrepancies in predicted 3D structures. Our analysis began with *altR*, a *tetR*-family regulator within *alt*-like gene clusters, revealing high structural similarities between BG and PA and BG and PTO, with zeal scores of 0.93 and 0.94, respectively. Further examination of secondary *altR* variants from PTO showed similar congruence, with scores ranging from 0.95 to 0.96. Extending our analysis to other genes such as *altA*, *altB*, *altC*, *altE*, and *altI*, we consistently observed high zeal scores (0.91 to 0.97) indicative of substantial structural similarity across different organisms. However, *altI* presented some structural discordance, with lower zeal scores down to 0.67, suggesting potential functional diversity. We expanded our study to include multiple sequence alignments of the five most frequently identified *alt*-like genes post-DeepBGC detection, followed by ITASSER-based 3D structural predictions. These comparisons involved a broad set of sequences, with resulting zeal scores ranging from 0.40 to 1.00, reflecting a wide diversity in structural similarity among the *altC* variants. Although the structural comparisons generally supported the structural resemblance across these genes, they did not provide a clear distinction between the datamined gene clusters. (figures 4 and 5; supplementary figures 1 and 2; supplementary files 1 and 2).

**Figure 3.**
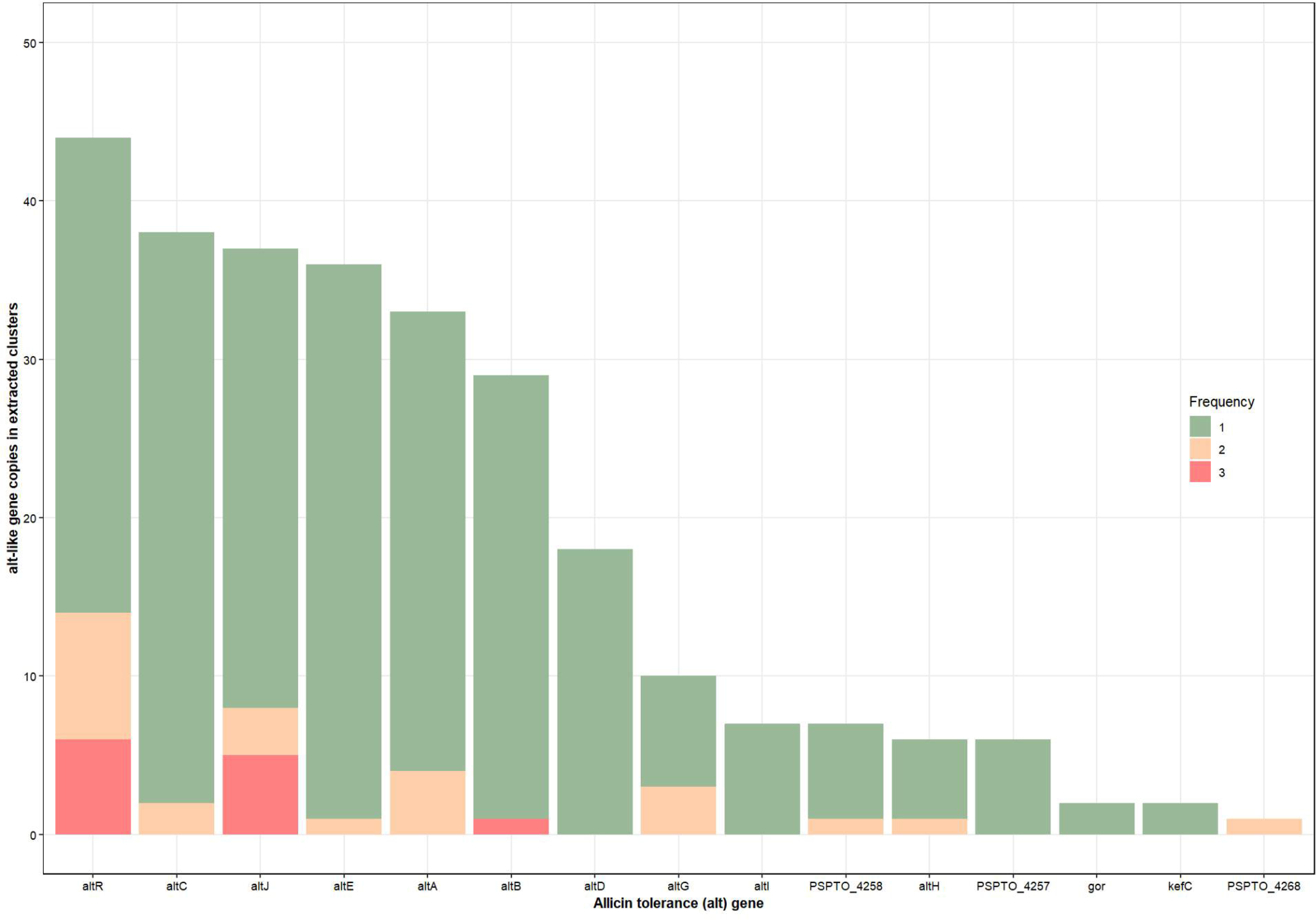
Frequency of allicin tolerance (*alt*)-like genes in DeepBGC extracted gene clusters from bacterial RefSeq database. Here, *alt*-like gene frequency was calculated from each of the representative gene clusters and added input clusters to determine the number of copies of alt-like genes present in each gene cluster. The graph is organized based on total gene count, with *altR* being the highest and *PSPTO_5268* the lowest. Green colors indicate the *alt*-like gene appeared only once in the extracted gene cluster. Yellow indicates the gene appeared twice in certain gene clusters. Pink indicates the gene appeared three times in certain gene clusters. Red indicates the gene appeared four times in certain gene clusters. *altC*, *altE*, *altA*, *altG*, *altH*, and *PSPTO_4258* appear once or twice in certain genomes. *PSPTO_5268* appeared twice in one extracted genome. *altI*, *altD*, *gor*, and *kefC* all appeared only once in their extracted genomes.

**Figure 4.**
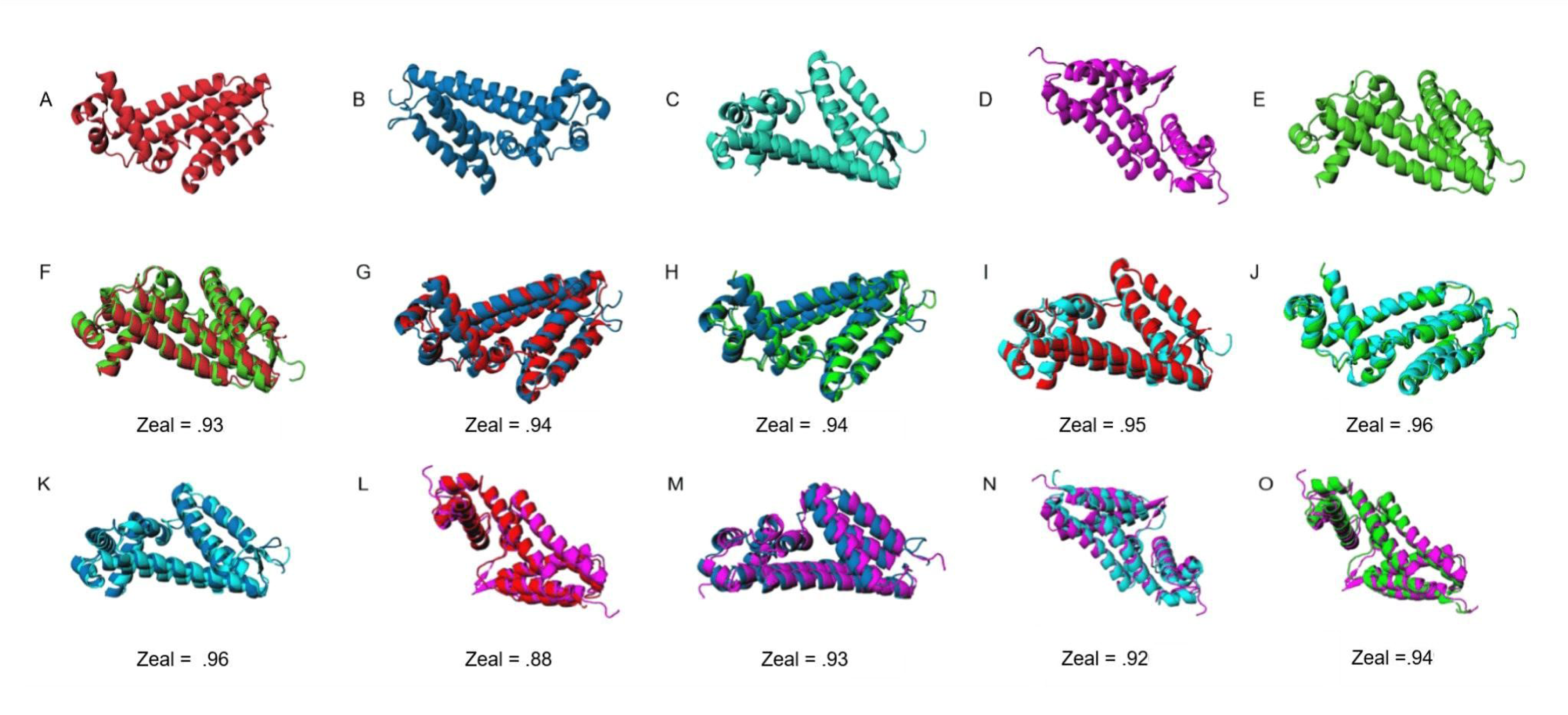
Comparative 3D superimposition of I-TASSER predicted *altR* repressors PNA 97-1R (PA), *Pseudomonas syringae* pv. tomato DC3000 (PTO), and an unrelated repressor from *Escherichia coli*, *nemR* (ECN). The initial row (A-E) represents all predicted protein structures used for downstream comparisons. The predicted model proteins are displayed as BG *altR* (A), PTO *altR* (B), PTO *out altR* (C), ECN (D), and PA *altR* (E). The 3D superimposition comparisons are shown in panels F-O. Values below each comparison refer to the Zeal score as predicted by the Zeal GUI (https://andrelab.lu.se/) and are an indication of shape similarity. For example, a zeal score of “1” indicates the same 3D protein shape. The “F” compares the predicted *altR* protein from BG vs. PA, while panels G and H represent the comparison of *altR* between BG vs. PTO and PA vs. PTO, respectively. The panels I, J, and K compare *altR* between BG, PA, and PTO vs. the PTO out_*altR* as indicated in Figure 1, respectively. The panels L, M, N, and O compare BG *altR*, PTO *altR*, PTO *out*_*altR*, and PA *altR* against ECN, respectively.

### Crosstree Comparisons between Protein Sequence Similarity and Gene Synteny Indicate vertical transmission and divergence of *alt* and *alt*-like Genes

To determine if there is any grouping of *alt*-like genes based on protein sequence similarity, we utilized RAxML to generate phylogenetic trees based on sequence similarity. Further, we used phytools to compare trees for pattern similarity. We utilized further R scripting to label the connecting lines with colors representing the terminal group these sequences belong to and their validation results. Bootstrap values for the trees appear low on several edges, indicating difficulty organizing groups effectively based on sequence. However, the comparison of the two trees together shows that the “core” *alt* proteins are primarily concordant with each other. While there are some potential notable exceptions, such as the *altC* from NZ_JACXQ010000006.1, the *Rahnella aqualitis* representative *alt*-like cluster, this appears to be due to the rotation of the tree as opposed to a biological reality. This opinion is further supported by both overlaying the validation data on these tree comparisons, where proteins with similar *alt* tolerance appear to be grouping together, and other comparison trees place the sequence much closer to the other gene synteny groups. These trees suggest that these collections of *alt*-like proteins appear to have independent evolutionary histories as vertically maintained genes despite being horizontally transferred. Further, the concordance of the validation data and these proteins seem to suggest specialization is occurring with the more robust *alt* phenotypes consistently grouping (supplementary folder 1).

### Phenotypic testing with Synthesized *altC*/*altE* Pairs Provides Evidence for Thiosulfinate Tolerance Functionality in Predicted *alt*-like Gene Clusters

To evaluate whether predicted *alt*-like clusters were legitimate and capable of conferring increased thiosulfinate tolerance phenotypes, we heterologously expressed synthesized *altC*/*altE* gene pairs representing key phylogenetic nodes in PA strain PNA 97-1R Δ*alt*, which lacks the functioning *alt* cluster and has poor thiosulfinate tolerance. Strains were grown in 50:50 LB onion extract as in Stice et al., and the mean area under the growth curve (AUC) was determined. Growth was compared against a thiosulfinate-sensitized PNA 97-1R Δ*alt* GFP expressing strain as a control and the PA wild-type strain (PNA 97-1R). Across all experiments, expression of GFP in PA PNA 97-1R Δ*alt* consistently showed the lowest growth of the inoculated OJ, indicating poor thiosulfinate tolerance.

In contrast, our positive control, PA PNA 97-1R WT, showed robust growth. Irrespective of *altC/altE* pairs from different bacteria, heterologous expression in PA PNA 97-1R Δ*alt* resulted in increased tolerance to thiosulfinate in our onion-juice (OJ) growth assay. The *altC/altE* pairs for bacteria that are closer to PA phylogenetically (*Erwinia persicina* CFBP8795, *Rahnella aquatillis* Ra9-2) tended to result in improved restoration of thiosulfinate tolerance to PNA 97-1R Δ*alt* compared with those that were phylogenetically distant (*Paenibacillus nuruki* TI45-13ar, *Burkholderia gladioli* BCC1802, and *Novosphingobium sp.* Chol11). However, an exception in this trend was observed with *Gluconobacter kondonii* (Dm-54). Despite its relative closedness with PA phylogenetically, the *altC/altE* heterologous expression in PA PNA 97-1R Δ*alt* did not result in consistent growth in the OJ growth assay, indicating comparatively lower tolerance to thiosulfinates (figure 6A-C and E). In addition, *Cronobacter dubliensis* (cro910B3) showed weaker tolerance but more robust tolerance than that of Gluconobacter kondonii (Dm-54) despite its relative closeness with PA phylogenetically. Overall, while all *altC/altE* pairs conferred increased thiosulfinate tolerance, the quantitative performance of individual *altC*/*altE* pairs is not easily predicted based solely on their phylogenetic similarity (figure 6D).

**Figure 5.**
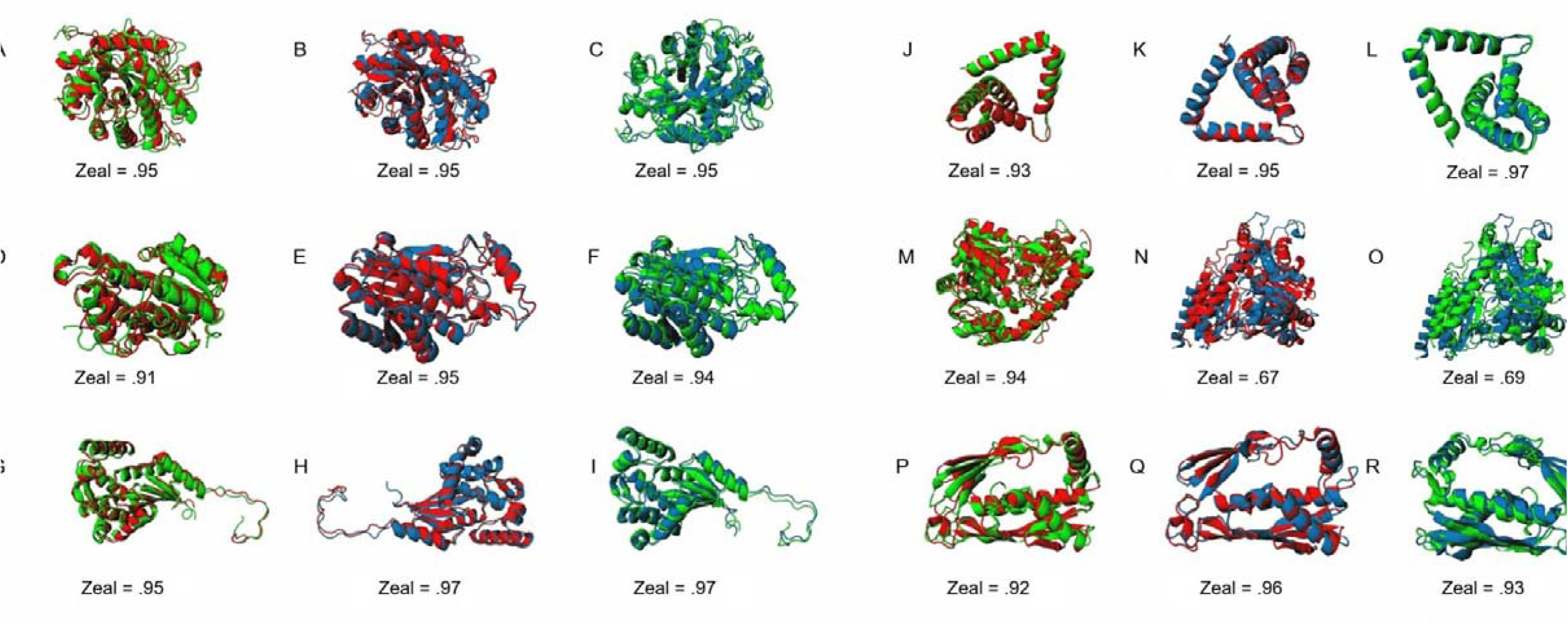
Comparative 3D superimposition of I-TASSER predicted allicin tolerance (*alt*) proteins between *Burkholderia gladioli* pv. *gladioli* FDAARGOS_389 (BG), *Pantoea ananatis* PNA 97-1R (PA), and *Pseudomonas syringae* pv. tomato DC3000 (PTO). BG proteins are colored red, PA proteins are colored green, and PTO proteins are colored blue for ease of visualization. Values below each comparison refer to the Zeal score as predicted by the Zeal GUI (https://andrelab.lu.se/) and are an indication of shape similarity. For example, a zeal score of “1” indicates the same shape. Each alt protein prediction is organized into three groups. The A, B, and C are comparisons of *altA* between BG vs. PA, BG vs. PTO, and PA vs. PTO, respectively. The panels D, E, and F are comparisons of *altB* between BG vs. PA, BG vs. PTO, and PA vs. PTO, respectively. The panels G, H, and I are comparisons of *altC* between BG vs. PA, BG vs. PTO, and PA vs. PTO, respectively. The panels J, K, and L are comparisons of *altE* between BG vs. PA, BG vs. PTO, and PA vs. PTO, respectively. The panels M, N, and O are comparisons of *altI* BG vs. PA, BG vs. PTO, and PA vs. PTO, respectively. The panels P, Q, and R are comparisons of *altJ* BG vs. PA, BG vs. PTO, and PA vs. PTO, respectively.

**Figure 6:**
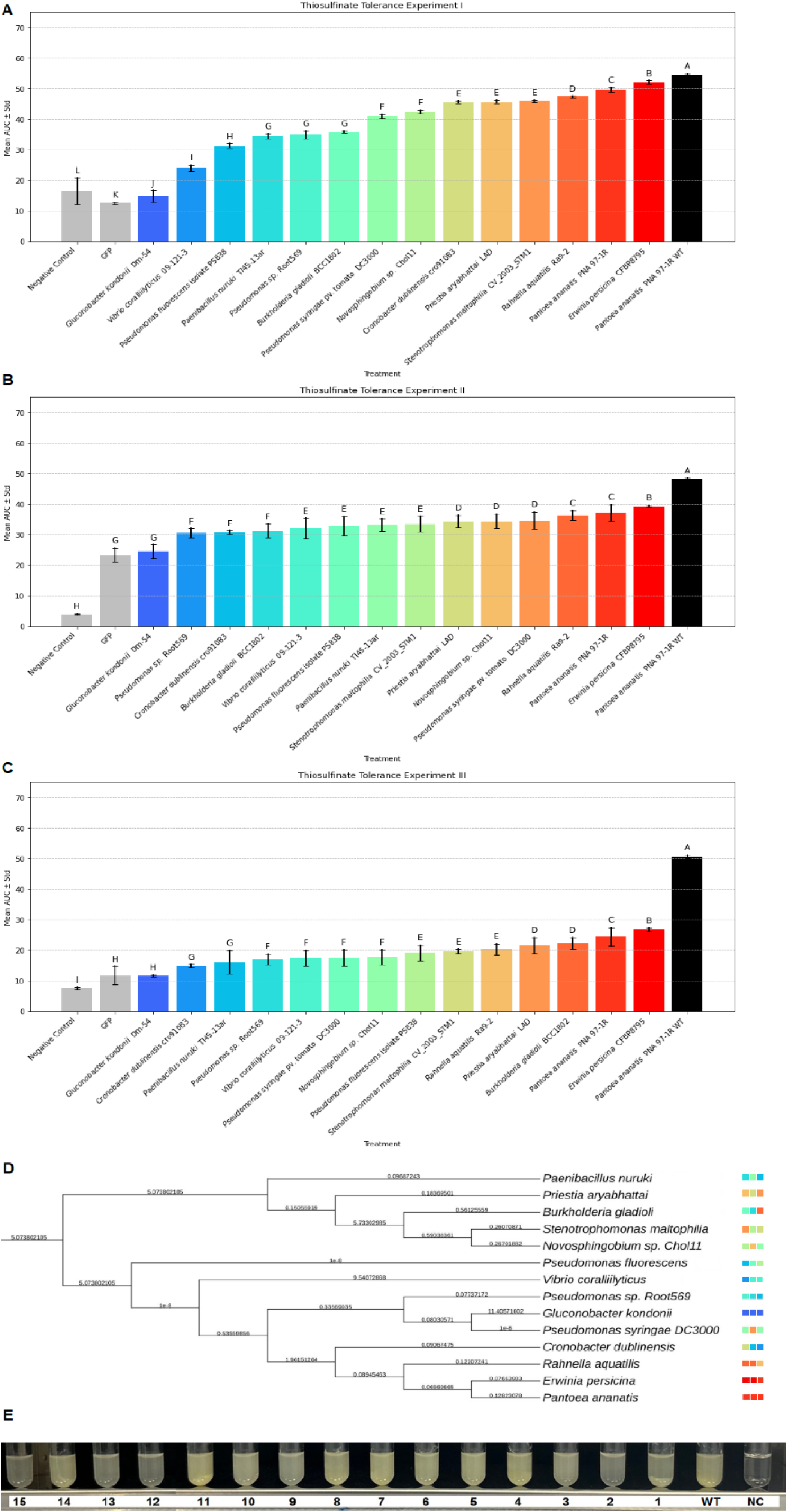
Thiosulfinate Tolerance was enhanced in *Pantoea ananatis* when transformed with *altC*/*altE* Pairs from diverse bacterial genera and their phylogenetic relationship among each other. Thiosulfinate tolerance of *Pantoea ananatis* PNA97-1 Δ*alt* was enhanced when transformed with *altC*/*altE* pairs representative of different bacterial genera and species conducted across three experiments, as well as their phylogenetic relationships. The top three bar charts in figure 6A, B, and C, represent three independent experiments, I, II, and III, respectively, with mean and standard error bars. The x-axis represents *P. ananatis* PNA 97-1R Δ*alt* transformed with *altC*/*altE* pairs from representative of different bacterial genera and species and controls (empty vector and water), while the y-axis shows the mean area under the curve (AUC) values. Statistical groupings are denoted by letters above the bars, with ‘A’ representing the group with the highest tolerance. Subsequent letters (B, C, D, E…) indicate progressively lower tolerance, based on Tukey’s HSD test results, with differences considered statistically significant at P<0.05. The negative control and GFP consistently show the lowest tolerance (labeled “E”), whereas *P. ananatis* PNA 97-1R WT and its variants exhibit the highest tolerance (labeled “A”). Figure 6D depicts the phylogenetic relationships among the tested *altC*/*altE* pairs in transformed *P. ananatis* from diverse bacterial genera and species, with branch lengths representing gene sequence distances. The color coding of the transformed *P. ananatis* strains with different *altC*/*altE* pairs matches the bars in the bar charts displayed above, providing a visual correlation between genetic similarity and thiosulfinate tolerance. Strain names were truncated for ease of visualization. For an alternative analysis of growth curve patterns across all experiments, please refer to the Euclidian distance tree in supplementary figure 3. Figure 6E shows a visual comparison of bacterial growth (in terms of turbidity) *P. ananatis* PNA 97-1R Δ*alt* transformed with *altC*/*altE* pairs from the following bacterial strains: 1: *Priestia aryabhattai* LAD; 2: *Novosphingobium* sp. Chol11; 3: GFP; 4: *Pseudomonas spp.* Root569; 5: *Stenotrophomonas maltophilia* CV_2003_STM1; 6: *Burkholderia gladioli* BCC1802; 7: *Pseudomonas fluorescens* PS838; 8: *Rahnella aquatilis* Ra9-2; 9: *Vibrio coralliilyticus* 09-121-3; 10: *P. ananatis* PNA 97-1 (WT, non-transformed); 11: *Pseudomonas syringae* DC3000; 12*: Paenibacillus nuruki* T145-13ar; 13: *Gluconobacter kondonii* Dm-54; 14: *Erwinia persicina* CFBP8795; 15*: Cronobacter dublinensis* cro91083, and NC: Negative control.

A hierarchical clustering analysis was conducted based on the Euclidean distance of the growth curves from the experiments to provide a comprehensive view of the growth response patterns across different bacterial strains. This analysis categorized the bacterial strains into clusters based on their growth response to thiosulfinate exposure. The hierarchical clustering dendrogram revealed distinct clusters, with each branch representing a similarity in growth responses among the strains. A distinct cluster formed by *G. kondonii* (Dm-54), *C. dublinensis* (cro910B3), and *P. nuruki* (TI45-13ar) indicates unique growth response profiles, which is supported by the unexpectedly poor performance of *G. kondonii* (Dm-54), and variability in responses from both *C. dublinensis* (cro910B3), and *P. nuruki* (TI45-13ar). A second significant cluster includes

*E. persicina* (CFBP8795) and *P. ananatis* 97-1R WT, showing more similar growth curves when compared to the remaining strains. These results are supported by the consistently high performance of the *E. persicina* (CFBP8795) *altC*/*altE* pair. The next group consists of similarly performing strains with *altC*/*altE* pairs from *Pseudomonas sp.* (Root569), *R. aquatilis* (Ra9-2), *Novosphingobium spp.* (Chol11), *P. aryabhattai* (LAD), and *S. maltophilia* (CV_2003_STM1) with *P. ananatis* (PNA 97-1R) placed in an intermediate rating with the previous group. These results are supported by the consistently high, but not as high, performance of *P. ananatis* (PNA 97-1R) when compared to *E. persicina* (CFBP8795), but not as variable as the remaining members of the group. V. coralliilyticus (09-121-3), and *B. gladioli* (BCC1802), *P. syringae* pv. tomato (DC3000), and *P. fluorescens* (PS838) are the final group. As expected, the GFP strain is positioned near the negative control, reinforcing its minimal growth and low tolerance to thiosulfinates because it lacked *alt* genes. The hierarchical clustering analysis provides a comprehensive view of the growth response patterns across different bacterial strains. It aligns with the tolerance experiments and phylogenetic analysis findings, demonstrating their similar growth profiles and tolerance mechanisms independently of protein sequence content or lineage.

### Binding Affinity Prediction with AI-BIND of *altR* Demonstrates NLP-like Techniques Are Effective for Predicting and Classifying *alt* and *alt*-like Proteins

To determine if NLP-like techniques for binding affinity prediction could be used to help differentiate between functional *alt* clusters and possible pseudo clusters, we utilized AI-Bind to screen our *altR* protein sequences against a library of small molecules collected from PubChem focusing on sulfur compounds (supplementary file 2). Due to the likelihood of noise among most of these binding predictions, rows within .001 similarity were extracted for individual assessment. Among the values extracted, several similarities among the columns can be seen, and these 28,481 predictions may be the primary drivers for the separations seen with the Levenshtein distance matrix. When the distance matrix is overlaid with the gene synteny plot and compared to the *altR* RAxML tree, it appears that the results generated from AI-Bind are capable of sorting *altR* proteins into their appropriate gene synteny groups. In addition, when integrating the findings with those from the experimental validation experiments, there is a pronounced division between *alt* clusters with robust phenotypes and those exhibiting weaker phenotypes. These results indicate that the results from AI-Bind could also sort *altR* proteins into groups that reflect the thiosulfinate tolerance of their respective *altC*/*altE* pair. As such, the binding predictions that AI-Bind produced may be helpful in further methodologies to automate the detection and distinction of *alt*, *alt*-like, and pseudo-*alt* proteins. This pattern further supports the notion that most, if not all, of the DeepBGC-identified representative *alt*-like clusters in this study are capable of functioning similarly to *alt*, a conclusion reinforced by the experimental validation results (figure 7).

**Figure 7:**
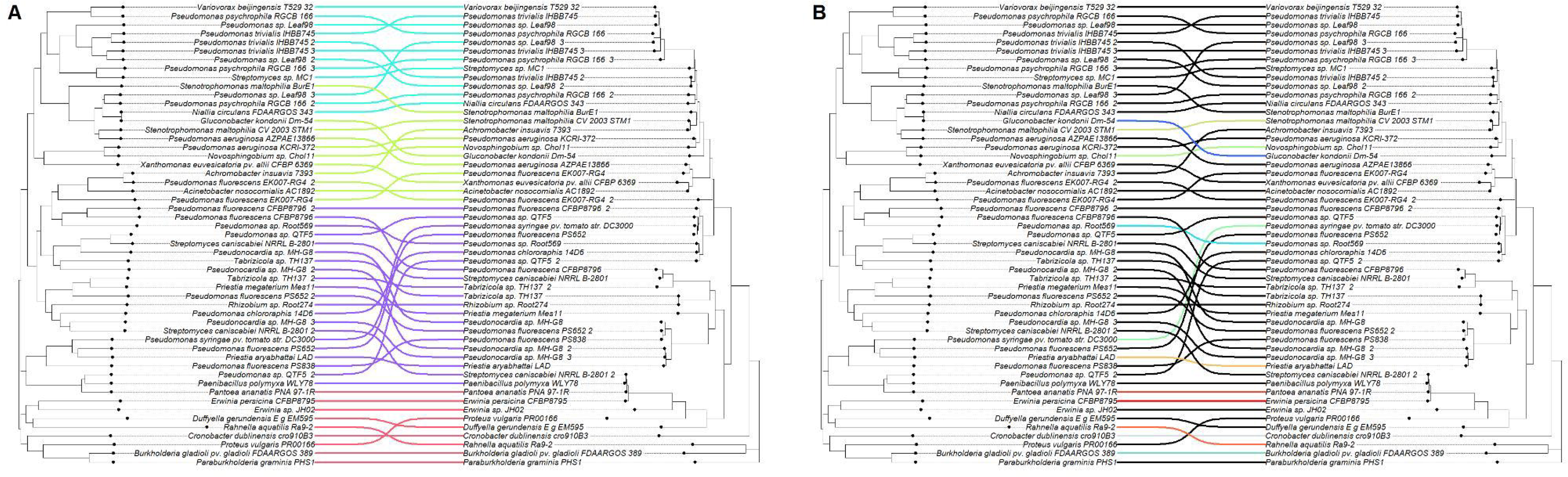
Comparison between gene synteny, protein similarity, and binding affinity prediction. Here are two schematic representations (A and B) that map the relationship between gene synteny alongside protein sequence similarity (Panel A) and between protein binding affinity predictions with corresponding protein sequences (Panel B). In Panel A, the branching lines are color-coded to distinguish between different gene synteny groups, which are identified as follows: pink for group I, green for group II, blue for group III, and purple for group IV, as previously defined in figure 2. Panel B contrasts the predicted binding affinities, as calculated by AI-Bind, of *altR* proteins against the similarity of the *altR* sequence. The coloration corresponds to the phenotypic data obtained from follow-up experimental validation, as described in figure 6, with the applied color serving as the “average” RGB value of the three colors.

## Discussion

Identifying *alt* and *alt*-like clusters poses challenges concerning variable gene synteny and divergent sequence similarities across bacterial genera. Specifically, *’alt* clusters’ refer to gene clusters experimentally validated to exhibit the thiosulfinate tolerance phenotype. Meanwhile, *’alt*-like clusters’ resemble these gene clusters in genetic composition but lack experimental validation for the associated phenotype. ‘Pseudo *alt* clusters,’ on the other hand, have been experimentally shown to not possess the phenotype despite their similarity in appearance to *alt* clusters. The three gene clusters utilized in our training set show little gene and protein sequence similarity and do not share overall gene synteny. Our analysis found a limited set of seven genes shared among our lab-validated gene clusters, with notable variations in gene and protein sequence similarities—highlighting the *altB* reductase in the SDR family oxidoreductase family as the most conserved element across these clusters. The observed sequence similarities range significantly, suggesting a nuanced spectrum of conservation and divergence within these gene clusters. Interestingly, despite the diversity in sequence similarity, the predicted protein structures demonstrated a surprising level of uniformity according to I-TASSER system evaluations. This uniformity, especially in the context of different pathogens from onion, underscores a potentially ancient divergence and pseudo-verticality for this horizontally transferred region.

Current *alt* clusters have been identified experimentally or predicted intuitively based on gene co-localization and annotation. However, this approach is difficult to rigorously codify and could lead to significant discrepancies between investigators. Further, we do not have a collection of pseudo*-alt* clusters to provide a comparison, exacerbating the difficulty in describing an actual *alt* cluster. Computational strategies, such as artificial intelligence methodologies like machine learning or deep learning, offer more sophisticated ways to “digitize” intuition for dissemination. In this work, we utilized an NLP-like method to generate a model capable of data mining these complex gene clusters with an unconventional training set of only 3 divergent validated gene clusters and, by extension, make a transition from bespoke manual curation of *alt* clusters into a streamlined process.

NLP in biology is becoming a valuable tool in studying gene function. There is a significant volume of genes that have an unknown function. By extension, we cannot access these genes’ full potential for biotechnology, agriculture, and medicine. In prokaryotes, for example, genes with a complementary function tend to group into biosynthetic gene clusters (14, 15, 16, 27). Relevant biosynthetic gene clusters can be detected and datamined by focusing on higher-order information and gene proximity. In this work, we used DeepBGC to train a model of our three previously validated *alt* clusters to overcome the limitations with more traditional sequence-only methods for data mining gene clusters, as well as explore the utilization of these techniques for identifying patterns that can be useful for identifying these clusters more robustly. DeepBGC utilizes Pfam information rather than the amino acid sequence to classify BGCs, with the additional caveat of understanding the importance of gene localization in gene clusters (24).

Utilizing vectorized Pfam domains and gene localization elegantly simulates our intuitive process to curate gene clusters and produces a tangible model that is more appropriate for rigorous scientific evaluation. Sequence-based methods, such as BLAST, had been insufficient for data mining these clusters across multiple genera due to low sequence similarity. However, the methodology utilized by DeepBGC produced a model that can successfully detect *alt* clusters reproducibly from diverse genera of bacteria, even breaking the Gram barrier. In an ideal scenario, a bioinformatician would have access to thousands of examples for their training set. In this study, we only had access to three validated examples of *alt*-clusters from PA, BG, and PTO. Despite this, we were able to successfully detect, retrieve, and validate several *alt* clusters that were previously undetectable. This methodology also alleviates the immense effort required to screen this expansive list of bacterial genomes. Utilizing NLP technologies in a biological context is a powerful tool for “standardizing” the intuitive extraction process.

We acknowledge that using the model provided in the supplementary materials, with a training set of only three clusters, is too broad to filter out background noise effectively. For example, our analysis incorrectly identified several gene clusters simply due to the presence of several tandem thioredoxin-related genes. Additionally, other clusters were mistakenly detected due to the presence of multiple copies of *tetR*-family repressor genes, leading to false classifications. Further, some datamined *alt* clusters would lose genes on the terminal ends of their gene cluster, but the same cluster from another genome would contain the entire expected sequence. This issue is resolved by running the model multiple times and determining the “average” cluster sequence. However, these types of errors are commented upon in the DeepBGC manuscript and are to be expected (24). As always, manual curation should be employed to ensure that AI models are behaving appropriately. Despite the occasional error in incredibly diverse genomes, when the model is run on the genome of an onion pathogen with a known *alt* cluster, deepBGC always performed the expected extraction.

We utilized another text-comparison technique to compare gene synteny. The complexity of the *alt* clusters often overwhelmed traditional DNA-sequence-based methods, frequently leading to system failures in organizing the information. However, by converting from one language to another and calculating the Levenshtein distance matrix, we successfully organized gene clusters into gene synteny groups quickly and reliably. The Levenshtein distance matrix is the “edit distance” between two strings. These are insertions, deletions, and substitutions (28, 29). We cut down the computational time and simplified the visualization process by converting our gene clusters into a color code and then a string representing these color codes. The application of language processing techniques is not limited to complex AI modeling or requires expensive computational equipment to be helpful. By applying the Levenshtein distance matrix, we can initiate the classification of *alt* clusters based on higher-order information, such as guilt-by-association syntax, in a comprehensive manner. Although the *alt* cluster exhibits many characteristics typical of horizontally transferred gene clusters, it appears to be maintained vertically within several bacterial genera. Despite this, gene synteny is not unique across bacterial genera, as specific gene patterns recur across multiple genera even if their sequence content is different.

When comparing the gene synteny to the experimental validation of *altC*/*altE* pairs, it appears that the *altC*/*altE* pairs from the first terminal group have more robust thiosulfinate tolerance restoration phenotypes than *altC*/*altE* pairs from other terminal groups. These *alt* clusters are also represented among many members of the enterobacteria; however, *alt*-like clusters from *Erwinia*/*Pantoea* displayed the strongest phenotype. A notable caveat with this methodology is that the strongest phenotype is observed when these clusters are expressed in *Pantoea*, potentially due to interactions and dependencies with other endogenous host factors. Based on the current information, gene cluster synteny alone is insufficient for comprehensively categorizing *alt*-like clusters. These results are unsurprising, as genes with distinct evolutionary histories can independently form gene clusters with similar synteny. However, repeating motifs among several gene clusters is a strong indication of collaboration for a phenotype, and we would argue that the guilt-by-association of these shared genes is still a substantial factor in identifying *alt*-like clusters, even if their motifs are not perfect indicators of *alt*-like phenotype performance in OJ.

In drug discovery, the conformation and 3D structure of molecules are critical, as small molecules must fit into a binding pocket of a target protein with a favorable reaction. Similarly, proteins that yield similar phenotypic functions are expected to have comparable shapes, regardless of their sequence similarity (30). This work explored the potential for predicted protein conformation to indicate the *alt* phenotype. The results of the *altC*/*altE* pair validation suggest that the 3D superimposition of our putative *alt*-like proteins to the *alt*-verified proteins may indicate qualitative but not quantitative phenotype. More proteomic experiments to compare proper *alt* and pseudo-*alt* proteins are necessary, as our validation experiment showed all selected *altC*/*altE* pairs provided thiosulfinate tolerance, with some exception to the pair derived from *Gluconobacter kondii* (Dm-54).

For example, when interpreting our phenotypic validation results in the context of our 3D superimposition, it is essential to note that the *altC*/*altE* pair from *Priestia aryabhattai* LAD (NZ_CP072478.1) exhibited a more robust thiosulfinate tolerance restoration phenotype compared to *Pseudomonas sp.* Root569 (NZ_LMGQ01000029.1), despite the latter having higher Zeal scores. This observation suggests that while structural similarities generally correlate with functional outcomes, exceptions highlight the complexity of phenotype-genotype relationships. Furthermore, *altC* variants with Zeal scores greater than 0.98 consistently supported more robust bacterial growth in our thiosulfinate tolerance growth assay, implying a potential threshold effect where high structural fidelity may enhance certain functional capabilities. Conversely, *altE* adds another layer of complexity; *Priestia aryabhattai* LAD (NZ_CP072478.1), with a lower Zeal score of only 0.69, showed a slightly stronger thiosulfinate tolerance restoration phenotype than *Novosphingobium sp*. Chol11 (NZ_OBMU01000004.1), which had a higher Zeal score of 0.94.

Additionally, our comparison of the *E. coli nemR* repressor with the four *altR* sequences in the genomes used for our training dataset revealed high similarity in their 3D protein structures. The *nemR* repressor in *E. coli* shows ranges from .88 to .94 similarity. In contrast, the other four exhibit similarities ranging from .92 to .98. This level of resemblance is expected, given that they all are annotated as *tetR* repressors. The *nemR* repressor in *E. coli* is assumed to be responsive to reactive chlorine (bleach) and nitrogen species [31]. As such, we find the protein shape to be helpful in providing a secondary opinion for the protein predictions, as apparent outliers can be screened independent of annotations but alone cannot be used to separate functional *alt* proteins from possible pseudo-*alt* proteins. These findings underscore the limitations of relying solely on structural predictions to infer functional characteristics, highlighting the need for more complex integrated approaches to classify *alt* and pseudo-*alt* proteins. This opinion is reinforced by the results of the AI-Bind screen, where we assessed whether screening for the potential binding affinity of proteins to a set of organo-sulfur molecules could help differentiate *alt*-like clusters.

We utilized AI-Bind to evaluate the predicted binding affinity of *altR* sequences against a library of 381,350 small molecules. We then calculated the string differences from a concatenation of the resulting scores to determine if the output could be informative for classification. Initially, the matrix generated from the AI-Bind average scores seemed discordant compared to the trees derived from protein sequence similarity. However, overlaying the AI-Bind prediction matrix with the data from the gene synteny matrix, as well as the result of the experimental validation, shows that binding affinity predictions from AI-Bind are capable of sorting *altR* proteins into groups that are reflective of our other screening methods, independently. These findings suggest that using average binding predictions may be an effective tool for further classifying *alt* clusters and separating *alt* and pseudo-*alt* proteins. It is important to note that AI-Bind, however, is not in and of itself utilized for the classification of proteins in this way and is only NLP-like in that it is performing its classification based on sequence data rather than utilizing more traditional NLP-like systems.

## Conclusions

NLP-like technologies are powerful tools to assist in the discovery and classification of gene clusters. Here, we generated a model capable of detecting and extracting *alt* clusters, validating the phenotype in transformed bacteria that previously lacked it. Despite its limited training set, the NLP-like algorithm used here demonstrated its capacity to identify several biologically relevant gene clusters. A model that quickly and accurately discovers and extracts *alt* clusters proves beneficial for diagnostic plant pathology and environmental bacteriology, particularly as the *alt* cluster is crucial for effectively colonizing *Allium* species or other thiosulfinate-producing hosts. The distribution of *alt* clusters beyond plant pathogens aligns with these secondary metabolites, shaping their microbial communities, as observed with the benzoxazinoid tolerance of maize root colonizers. Employing sophisticated NLP-like tools may revolutionize our understanding of critical gene clusters that facilitate complex host-microbe interactions, potentially leading to breakthroughs in several multidisciplinary fields. In developing a more robust *alt* cluster detection system, integrating models that encompass Pfam domains, gene localization, and predicted binding affinity might be sufficient to distinguish between *alt* clusters—those experimentally validated to function—and pseudo *alt* clusters, which appear similar but are experimentally validated not to possess the phenotype.

## Materials and Methods

### *alt* Gene Cluster Seed Sequences

For this work, we used three validated *alt* clusters for DeepBGC training. Each cluster is distinct in both gene sequence and gene synteny. The 11-gene *Pantoea alt* cluster was used from *Pantoea ananatis* strain PNA 97-1R plasmid unamed2 (NCBI accession: PRJNA384061). The 7-gene *Burkholderia alt* cluster was used from *Burkholderia gladioli* pv. *gladioli* strain FDAARGOS_389, plasmid unnamed (NCBI accession: PRJNA231221). The *Pseudomonas alt* cluster was used from *Pseudomonas syringae* pv. *tomato* str. DC3000, complete genome (NCBI accession: PRJNA57967) (supplementary folder 2).

### Gene/Protein sequence comparisons for validated *alt* clusters

To understand sequence similarity between validated *alt* genes, we performed multiple sequence alignments of protein and nucleic acid sequences at default settings using the Clustal Omega online server (32).

### DeepBGC training and RefSeq screening

To determine if AI trained on higher-order information can assist in efficiently datamining *alt*-like clusters from a collection of genomes, we trained the DeepBGC model on our small sample size of 3 validated *alt* clusters. The *alt* detection model was trained using the author’s supplied negative dataset “GeneSwap_Negatives.pfam.tsv” and ran with the default provided “deepbgc.json” with DeepBGC version 0.1.27. DeepBGC training on the initial three *alt* sequences was repeated 15 times and the reports were compared to assess model performance. DeepBGC options on the database data mining included a minimum protein count of 4 and a minimum score of .9. RefSeq genomes were separated into 48 sub-directories of 5,000 genomes, and DeepBGC jobs were submitted to the UGA GACRC via an array element on the batch partition. We scanned 238,362 genomes using this model from the NCBI bacterial refseq database. The genomes were downloaded via the NCBI FTP service, and the assembly list is provided (supplementary files 3 and 4) (24).

### Filtering gene cluster representation via MMseqs2

To compress the DeepBGC extractions into a smaller representation for analysis, we used MMseqs2 release 13-45111 to generate representative sequences. The options used were a query coverage of 90%, sequence homology of 75%, and connected component clustering. These options allow for a “core” representative sequence with leniency for small changes in gene presence or absence (33).

### BLAST of NCBI GenBank for Representative Sequence Diversity

The final selection of 47 representative *alt*-like clusters was utilized as the query sequence for both the collection of putative *alt*-like gene clusters from DeepBGC and NCBI GenBank, following similar rules to the MMSEQ2 redundancy filtering. Recovered species were then counted and organized into a list for the calculation of the Shannon-Wiener Index via our own Python script.

### Gene synteny comparisons

During our manual curations of *alt*-like clusters we noticed a pattern where gene synteny was conserved among bacterial genera. For those clusters, we generated cluster comparisons with sequence data, as the method for data mining was determinate upon them. However, post-DeepBGC screening of RefSeq, we found many *alt*-like clusters that share low enough sequence similarity between genes of similar annotation that several methods for cluster comparison would fail. To overcome this barrier, we assigned a color code to *alt*-like genes that received pfam tags like our test run of the original test sequences. To optimize the human capacity to read the information and remove unintended bias between colors, we assigned several shades of green to *alt*-like genes and grey color to genes that are not relevant. We then used an Excel script to convert these colors into color codes. These codes were then concatenated into strings and underwent a Levenstein distance matrix calculation using the Levenstein and Dendropy Python packages [34, 35]. After initial tree construction, further manual curation was applied to finalize *alt*-like representatives by selecting gene clusters with at least 3 unique *alt*-like pfam tags that match those applied to the seed clusters and the removal of clusters that were split into separate contigs.

### Generating 3D protein models and Zeal score comparisons

While the previous methods make gene comparisons primarily on sequence or trained guilt-by-association with higher-order information, we wanted to compare *alt*-like proteins for potential structure diversity or abnormalities directly. Models for select *alt*-like proteins were generated using I-TASSER 5.2 with the -LBS option set to true. Predicted protein 3D models were then compared using the Zeal GUI with global alignment. (36, 37).

### Protein-ligand binding prediction

The *alt*, and *alt*-like mechanisms of action are currently unknown. However, it is reasonable to suspect that the binding interactions between chemicals and proteins would be essential in defining *alt*, *alt*-like, and pseudo-*alt* gene cluster classes. Due to the computationally expensive nature of drug-target binding predictions we turned to using AI-Bind, a deep-neural network designed for a more generalizable prediction of binding between proteins and small molecules. A comprehensive list of small molecule SMILES and InChiKeys were downloaded from the PubChem database with the following search terms: “allyl, cysteine sulfoxide, disulfide, polysulfide, S-Nitrosothiol, sulfenic acid, sulfenic, Sulfimide, sulfinic acid, silfinic, sulfone, Sulfonic acid, Sulfonic, Sulfonium, sulfoxide, sulfoximide, Sulfurane, thiolaldehyde, thioamide, thiocarbonyl, thiocarboxylic acid, thioester, thio, thiosulfinate, 316263-glutamylcysteine, and s-Allylmercaptoglutathione.” The results were concatenated, and duplicate entries were removed for a final list size of 381,349 small molecules. In our final representative dataset, these were then screened against the 53 *altR*-like proteins that received Pfam tags from deepBGC. Binding results from the *altR*-like proteins were converted into strings and compared via the calculation of the Levenstein distance matrix above to produce a neighbor-joining tree for ease of comparison (38).

### *altC*/*altE* Validation

To validate the representative *alt*-like clusters produced by DeepBGC, we conducted an onion-juice (OJ) growth assay. Previous research has demonstrated that the presence of *alt*C alone is sufficient to determine an *alt* phenotype. As such, we selected 14 *altC* genes (supplementary table 2) for validation, along with their potential *altE* partner if present. These 14 gene pairs (*altC*/*altE*) were inserted into Twist Bioscience’s pENTR plasmid and inserted into *P. ananatis* PNA 97-1^R^ Δ*alt* [10] by the following method.

### Electroporation and Confirmation

The recipient strain’s electrocompetent (e-comp) cells (*P. ananatis* PNA 97-1R *WT* and Δ*alt*) were prepared using standard methods. Plasmid constructs were electroporated into the recipient cells at 1.8kV. Transformed cells were mixed in 1 ml LB and left for incubation at 28°C for an hour. Post-incubation, cells were pelleted, resuspended in LB, and plated onto LB+Km plates. Individual transformed recipient cell colonies were grown overnight in LB+Km broth. Plasmids were extracted and sequenced to confirm the insertion of the pENTR plasmid constructs into recipient cells.

The plasmid pENTR::GFP served as an empty vector and was inserted into both PNA 97-1^R^ *WT* and Δ*alt* strains, which acted as positive and negative controls for the onion-juice growth assay. The remaining 14 plasmid inserts were transformed into PNA 97-1^R^ Δ*alt* strains.

### Preparation of Onion Juice Extract

Onion juice was extracted using Juicer method (10). One yellow onion bulb (400-500 g) was processed through an industrial strength juicer, resulting in 300-400 mL of crude onion extract. The extract was then centrifuged at 10,000 g for 1.5 hours at 4°C. After centrifugation, the supernatant was carefully removed and filtered through a Nalgene disposable vacuum filter sterilization unit. The onion juice was then stored at −20°C for future use.

### Liquid Growth Assay

The growth assay used 100-well honeycomb plates with the BioScreen C system (Lab Systems Helsinki, Finland). Seven-day-old OJ was utilized for the assay. 16 bacterial strains culture was started on LB+Km plates, and overnight cultures were prepared the following day in LB+Km broth from single colonies. On the third day, the growth assay was conducted for 48 hours with low agitation at 28°C. The growth media consisted of LB supplemented with an equal volume of onion juice. The experiment included 16 test strains (figure 6), including PNA 97-1 WT pENTR::GFP and Δ*alt* pENTR::GFP strains, and a negative control (LB+OJ). Each well contained 400 µL of a mixture comprising 360 µL of growth media (LB+OJ) and 40 µL of a bacterial suspension with an OD600 of 0.5 in sterile dH2O, with a minimum of 5 well replicates. Absorbance values were measured every 30 minutes for 48 hours. Each experimental assay was repeated twice for biological replication.

### Euclidean Distance Comparison of Treatment Groups

CSV files generated from three growth phases from our experiment, the lag, log, and stationary phases, were used to compare Euclidean distances between the growth curves of each treatment group via a Python script. For each phase, mean trendlines were calculated by averaging the sample data for each treatment group. If trendlines varied in length, shorter sequences were padded with NaN values to match the most extended sequence in each phase. Euclidean distance matrices were generated using the pdist function from the scipy.spatial.distance module, with NaN values imputed using the SimpleImputer with a mean strategy. For each phase, pairwise distances were calculated between all samples, and the resulting distance matrices were saved as CSV files. The average distances within and between treatment groups were computed to create symmetric group-level distance matrices that represented the average pairwise Euclidean distances for each treatment group in each phase. Hierarchical clustering was performed on these group-level matrices using the linkage method with ‘average’ linkage, and dendrograms were generated to visualize the clustering relationships between treatment groups. Dendrogram branches were color-coded according to predefined treatment groups to reflect their clustering pattern. The aggregated group average distances across all phases were calculated to compare the treatment groups’ growth patterns. A neighbor-joining tree for the combined dataset was generated and saved for visualizing the clustering results (supplementary figure 3).

### Tree Comparisons

Maximum likelihood trees were compared against each other using the phytools R package (39). For figure 7, edges between the nodes were organized based on their color code.

## Supporting information

All_Supplemental_Files

## Acknowledgments

We acknowledge Ms. Kailey Hara for her assistance with reformatting figure 1A with her expertise as a digital artist. This study was supported in part by resources and technical expertise from the Georgia Advanced Computing Resource Center, a partnership between the University of Georgia Office of the Vice President for Research and the Office of the Vice President for Information Technology.

