## Supplementary material for "Natural Language Processing-like Deep Learning Aided in Identification and Validation of Thiosulfinate Tolerance Clusters in Diverse Bacteria": All_Supplemental_Files: Supplementary legends.docx

**Supplementary file 1. Comparative zeal scores of various protein sequences.** Zeal scores for protein sequences from various bacterial genomes, evaluating their 3D structural superimposition across different bacterial genera. The zeal score is a metric used to assess the similarity of protein shapes, which is crucial for understanding functional similarities, especially in proteins with low sequence homology but similar surface structures. The columns include: Sequence (identifier of the protein sequence with accession numbers), *Pantoea* Zeal Score, *Burkholderia* Zeal Score, *Pseudomonas* Zeal Score, and *Pseudomonas* 2 Zeal Score. Each row corresponds to a specific protein sequence and its associated zeal scores, providing a comparative analysis of protein 3D structural alignment in *Pantoea*, *Burkholderia*, and two strains of *Pseudomonas* compared to other *alt*-like proteins ([Oxford Academic](https://academic.oup.com/bioinformatics/article/37/18/2874/6194581#:~:text=URL%3A%20https%3A%2F%2Facademic.oup.com%2Fbioinformatics%2Farticle%2F37%2F18%2F2874%2F6194581%0ALoading...%0AVisible%3A%200%25%20)) ([AndreLab](https://andrelab.lu.se/index.php/research/projects/35-zeal-protein-structure-alignment-based-on-shape-similar" \t "_blank)).

**Supplementary file 2. Comparative predictions from AiBind for several altR repressors**. The large Ai-BIND output file shows the average predictions of binding affinities across various bacterial strains using the AI-Bind algorithm. AI-Bind predicts binding affinities based on structural and sequence data, providing insights into the interactions between repressors and their targets, here a library of all results using these search terms from pubchem: “allyl, cysteine sulfoxide, disulfide, polysulfide, S-Nitrosothiol, sulfenic acid, sulfenic, Sulfimide, sulfinic acid, silfinic, sulfone, Sulfonic acid, Sulfonic, Sulfonium, sulfoxide, sulfoximide, Sulfurane, thiolaldehyde, thioamide, thiocarbonyl, thiocarboxylic acid, thioester, thio, thiosulfinate, 316263-glutamylcysteine, and s-Allylmercaptoglutathione, which indicate differences in binding affinity between *altR* repressors that may be useful in *alt* classification.

**Supplementary file 3. List of accessions downloaded from NCBI RefSeq and utilized for genome data mining**. This file contains a comprehensive list of accession numbers from the NCBI RefSeq database, which were used to obtain genomic sequences for mining allicin tolerance gene clusters. The accession numbers represent a variety of bacterial strains that were analyzed using the DeepBGC machine learning model to identify biosynthetic gene clusters (BGCs) associated with allicin tolerance (*alt*).

**Supplementary file 4.** **Detection of allicin tolerance gene clusters using a machine learning model from the DeepBGC package**. The provided file, Alt_like_BGC_Detector.pkl, contains a trained machine learning model designed to identify allicin tolerance (*alt*) gene clusters (BGCs) within genomic data. DeepBGC is a Python package that facilitates the identification and analysis of biosynthetic gene clusters using advanced machine learning techniques. This model leverages sequence features and patterns specific to allicin tolerance to predict the presence of relevant gene clusters in various bacterial genomes.

**Supplementary figure 1**. Comparative 3D superimposition of I-TASSER predicted allicin tolerance (*alt*) repressors between *Burkholderia gladioli* pv*. gladioli* FDAARGOS_389 (BG), *Pantoea ananatis* PNA 97-1R (PA), and *Pseudomonas syringae* pv. tomato DC3000 (PTO) continuously rotating. The initial row (A-E) represents all predicted protein structures used for downstream comparisons. The predicted model proteins are displayed as BG *altR* (A), PTO *altR* (B), PTO *out*_*altR* (C), PTO *PSPTO_4268* (D), and PA *altR* (E). A sulfate permease family protein model was used as a discordant/negative control to display a discordant protein superimposition. The sulfate permease family protein, *PSPTO_4268* (PTOS) (D) contains an authentic frame shift that is not the result of a sequencing artifact; the gene is disrupted by an insertion sequence. The 3D superimposition comparisons are shown in panels F-O. Values below each comparison refers to the Zeal score as predicted by the Zeal GUI (https://andrelab.lu.se/) and is an indication of shape similarity. For example, a Zeal score of “1” indicates the same 3D protein shape. The “F” compares the predicted *altR* protein from BG vs. PA while panels G and H represent the comparison of *altR* between BG vs PTO and PA vs PTO, respectively. The panels I, J, and K compare *altR* between BG, PA, and PTO vs the PTO out_*altR* as indicated in Figure 1, respectively. L, M, N, and O compare BG *altR*, PTO *altR*, PTO *out*_*altR*, and PA *altR* against PTO PTOS, respectively. Protein models were set to spin at an x-axis setting of 0, a z-axis of 0, and a y-axis of 20 to allow for a more holistic view of protein differences in 3D space using the jsmol option with the Zeal GUI.

**Supplementary figure 2.** Comparative 3D superimposition of I-TASSER predicted allicin tolerance (*alt*) proteins between *Burkholderia gladioli* pv. *gladioli* FDAARGOS_389 (BG), *Pantoea ananatis* PNA 97-1R (PA), and *Pseudomonas syringae* pv. tomato DC3000 (PTO) constantly rotating. BG proteins are colored red, PA proteins are colored green, and PTO proteins are colored blue for ease of visualization. Values below each comparison refers to the Zeal score as predicted by the Zeal GUI (https://andrelab.lu.se/) and is an indication of shape similarity. For example, a Zeal score of “1” indicates the same shape. Each alt protein prediction is organized into groups of three’s. A, B, and C are comparisons of *altA* between BG vs. PA, BG vs.PTO, and PA vs.PTO, respectively. The D, E, and F are comparisons of *altB* between BG vs.PA, BG vs. PTO, and PA vs. PTO, respectively. The G, H, and I are comparisons of *altC* between BG vs. PA, BG vs. PTO, and PA vs. PTO, respectively. The J, K, and L are comparisons of *altE* between BG vs. PA, BG vs. PTO, and PA vs. PTO, respectively. The M, N, and O are comparisons of *altI* BG vs. PA, BG vs. PTO, and PA vs. PTO, respectively. The P, Q and R are comparisons of *altJ* BG vs. PA, BG vs. PTO, and PA vs. PTO, respectively. Protein models were set to spin at an x-axis setting of 0, a z-axis of 0, and a y-axis of 20 to allow for a more holistic view of protein differences in 3D space using the jsmol option with the Zeal GUI.

**Supplementary table 1.** A detailed summary of the BLAST results for various bacterial sequences analyzed using DeepBGC and GenBank databases. Each sequence is identified by its unique ID, including accession numbers. The table includes the number of hits recovered by DeepBGC, the dominant genus or species identified, and the percentage of this dominant species, if applicable. Additionally, the Shannon-Wiener Index is provided to indicate the diversity within the environment. The table also includes the number of BLAST hits from GenBank, the dominant genus or species identified from these hits, the percentage of this species, if applicable, and another Shannon-Wiener Index value based on GenBank results. This supplementary data supports the main findings by detailing the diversity and dominance of bacterial species identified through BLAST analysis, highlighting the microbial composition and potential environmental impacts.

**Supplementary table 2**. This table contains the sequence information of the *altC*/*altE* pairs used for the phenotype validation.

**Supplementary folder 1**. A collection of images showcasing that gene synteny among the *alt*-like gene clusters is maintained across *alt*-like protein sequences despite the atypical nature of the *alt* gene cluster. These *alt* cluster sequences have a wide range of synteny and overall sequence similarity, and *alt* genes do not need to cohabitate as the genes are not collaboratively synthesizing a molecule for the phenotype.

**Supplementary folder 2**. This folder contains the GFF files used as the input sequences for DeepBGC.
