## Supplementary figures and images for "Natural Language Processing-like Deep Learning Aided in Identification and Validation of Thiosulfinate Tolerance Clusters in Diverse Bacteria"

### altCvsaltA.png

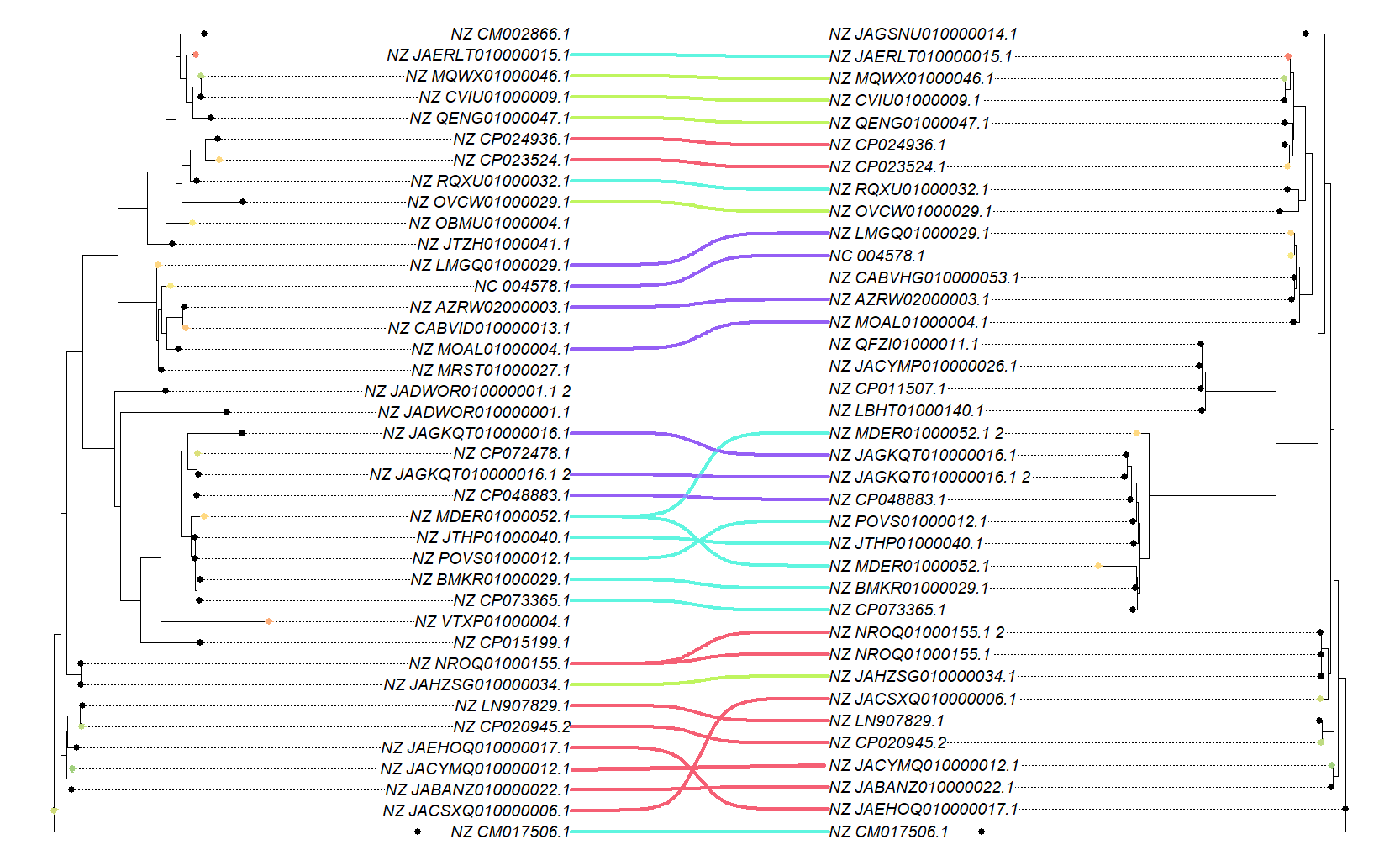

### altCvsaltE.png

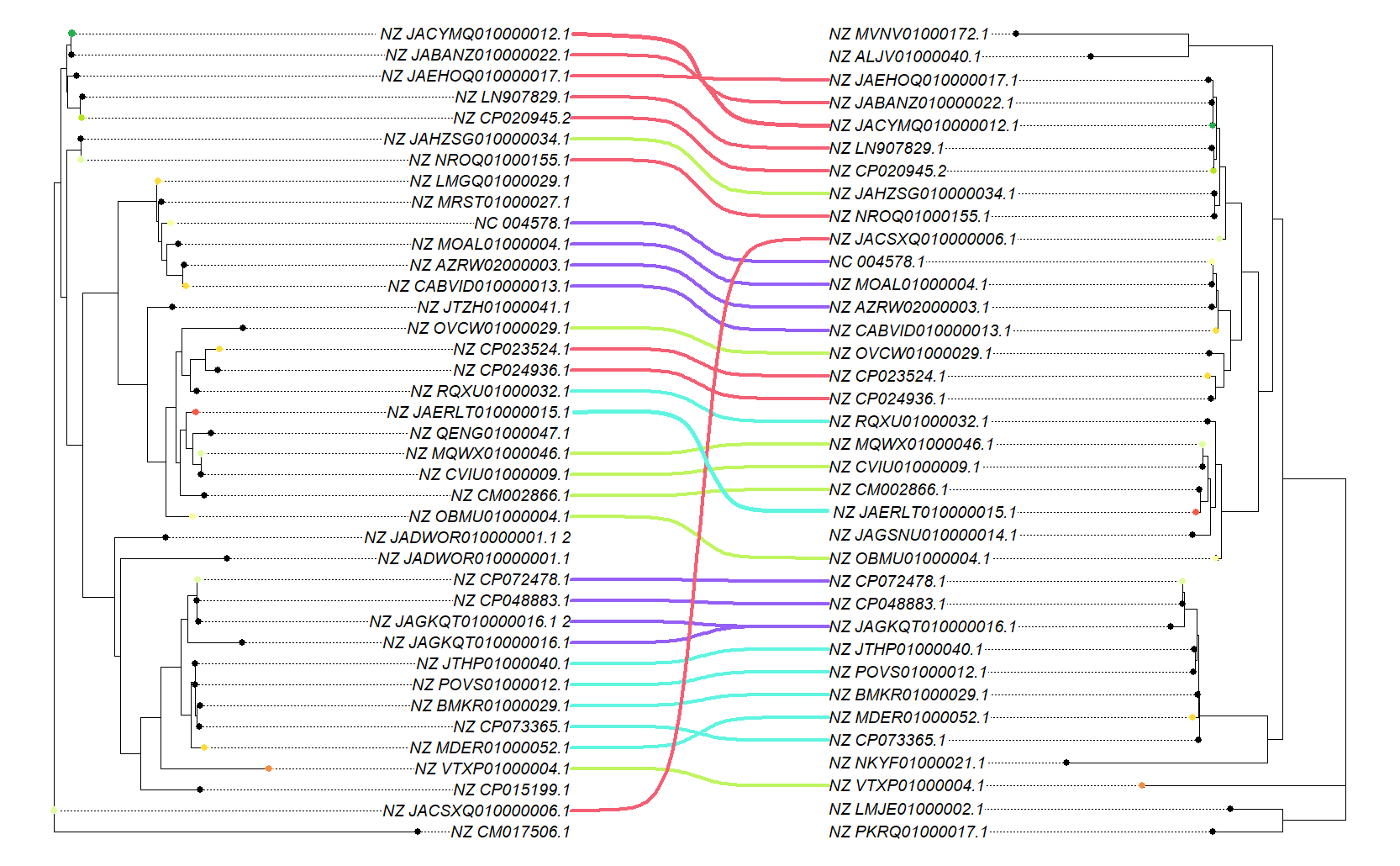

### altCvsaltJ.png

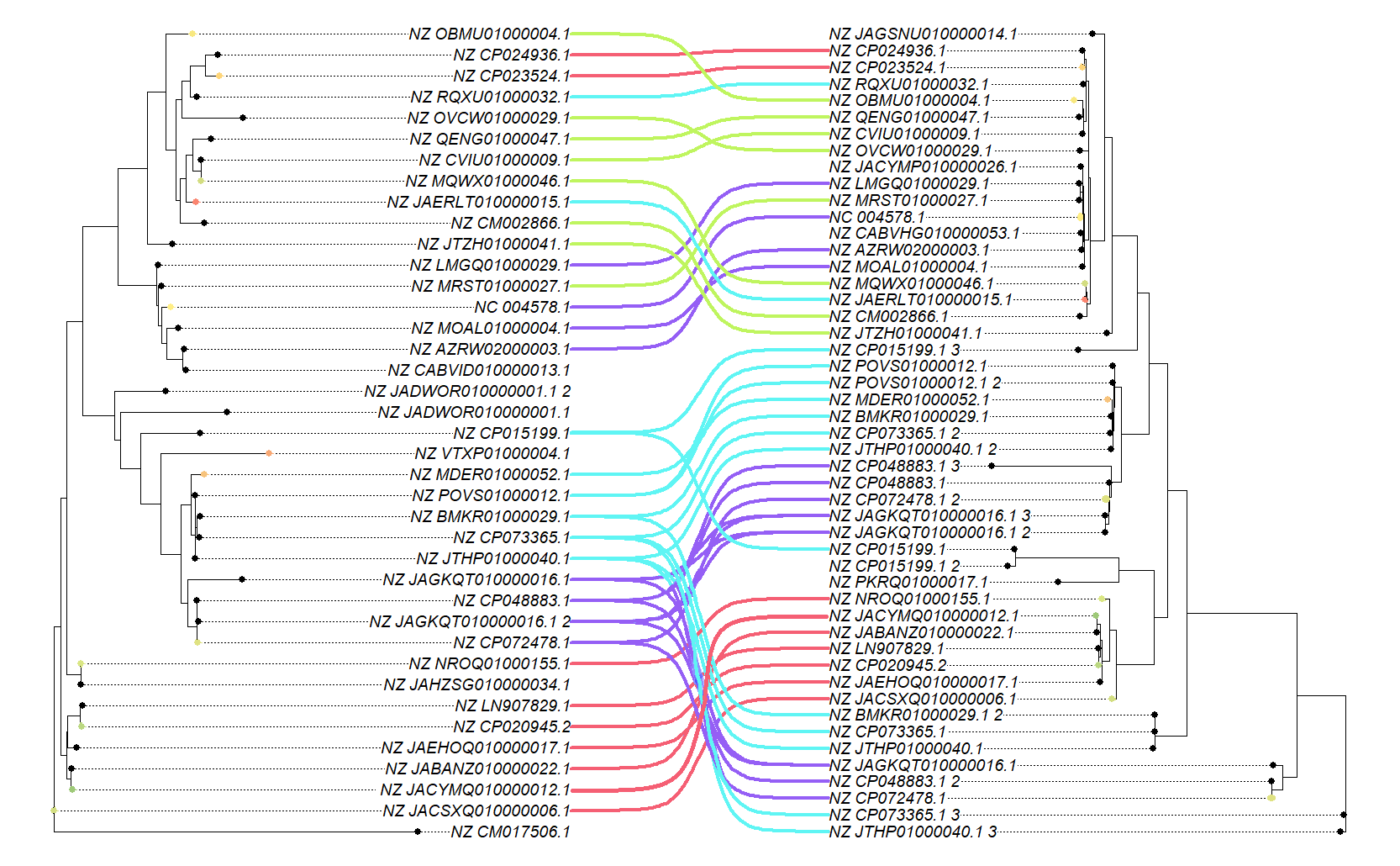

### altCvsaltR.png

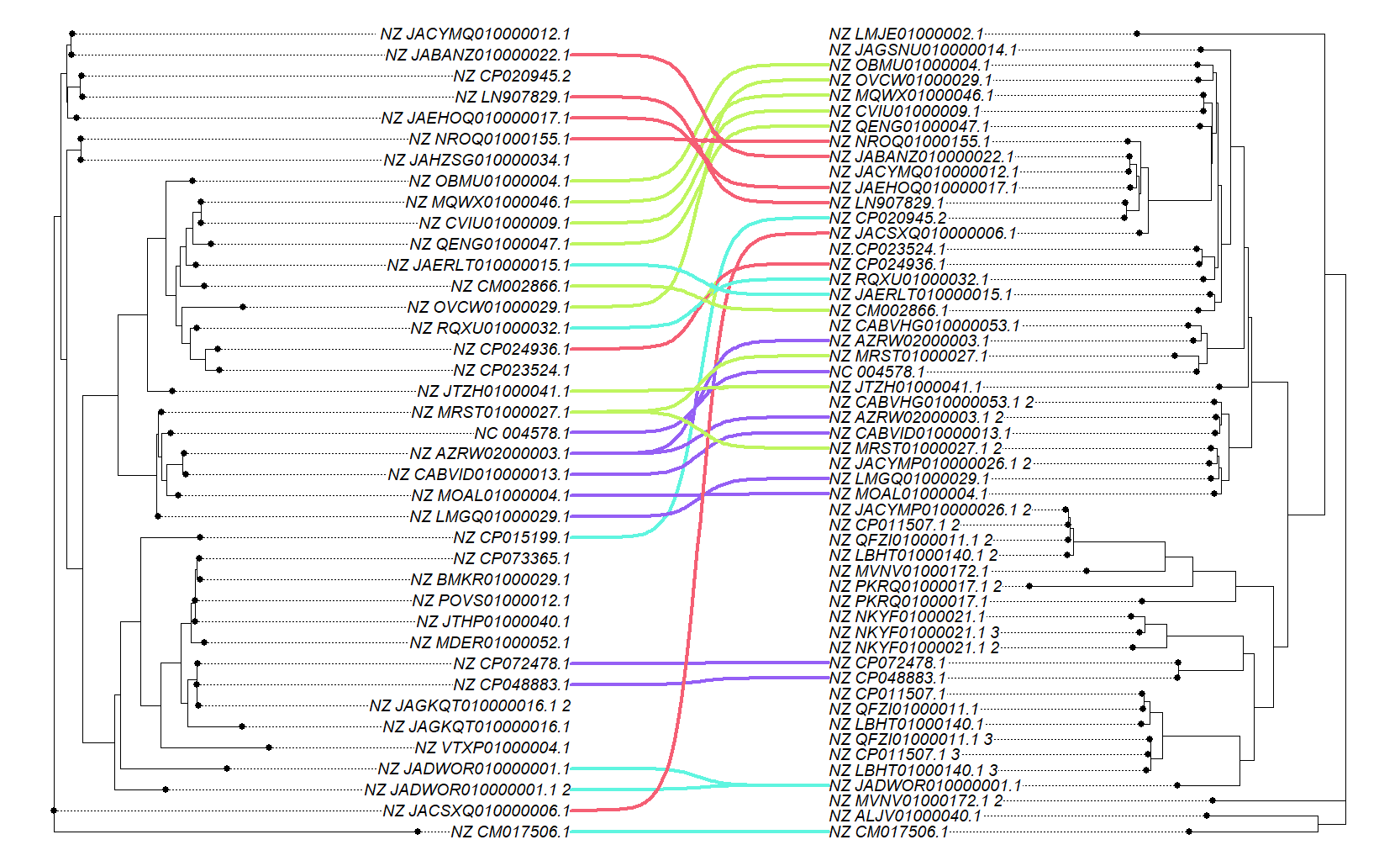

### altEvsaltA.png

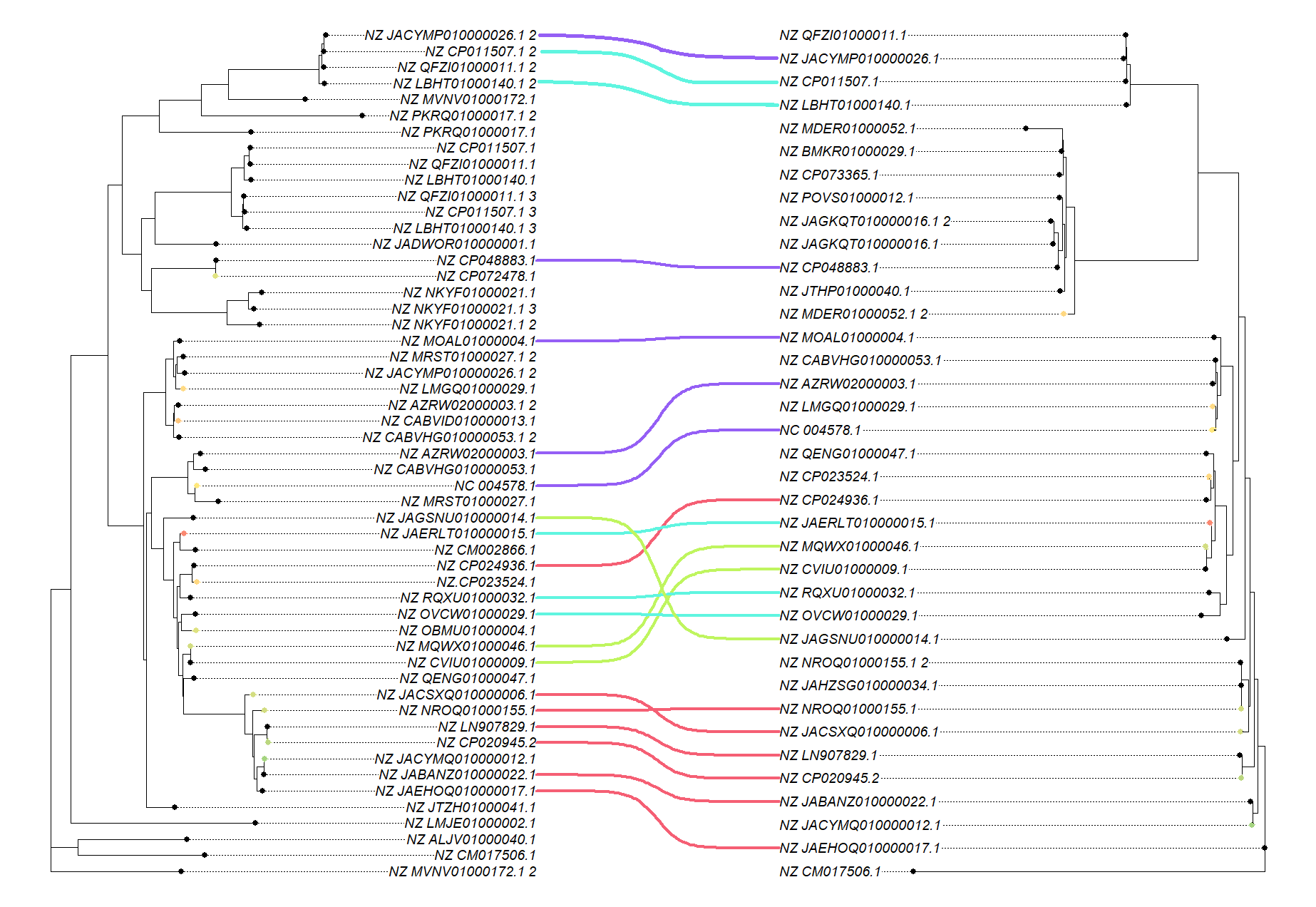

### altEvsaltJ.png

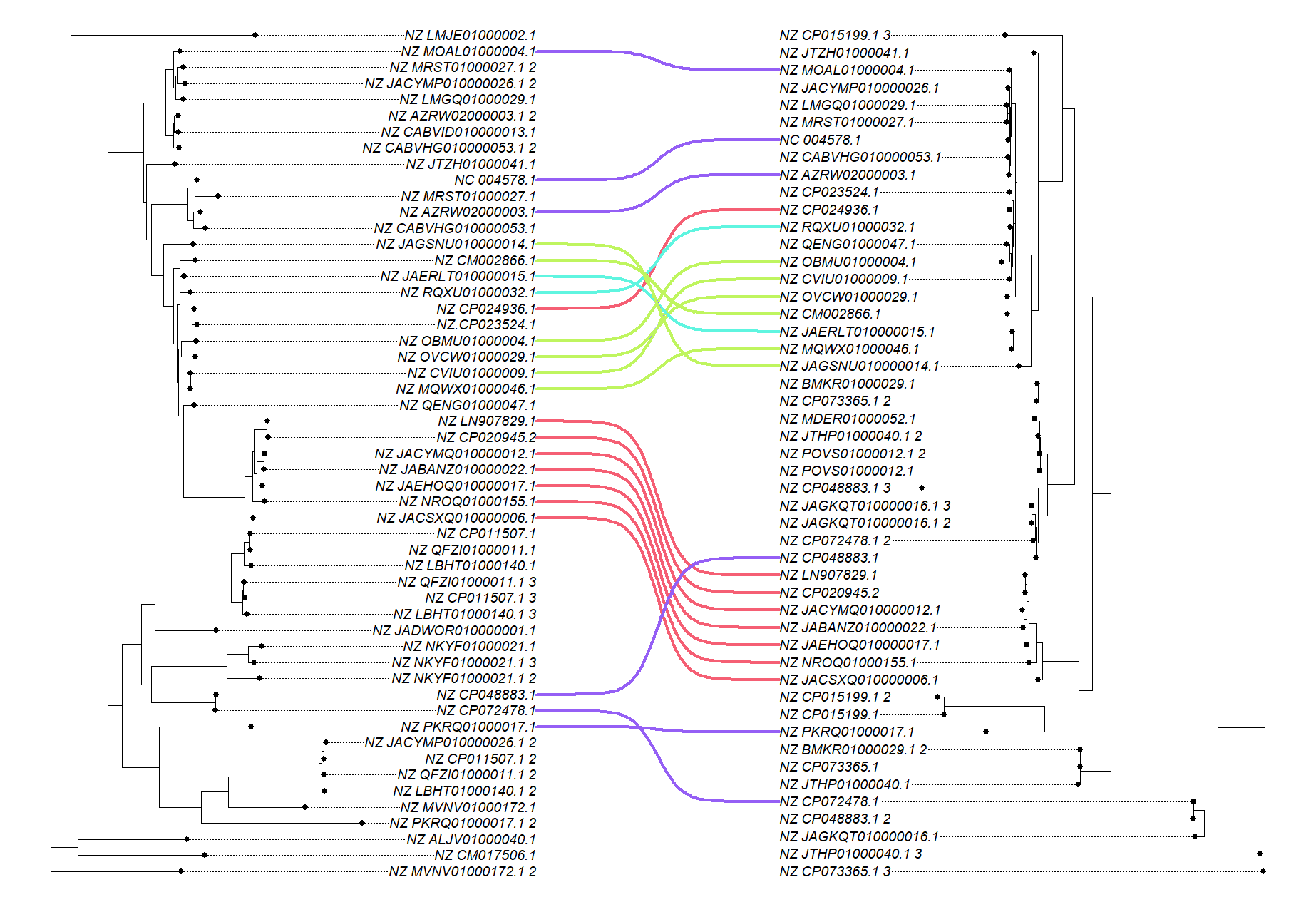

### altJvsaltA.png

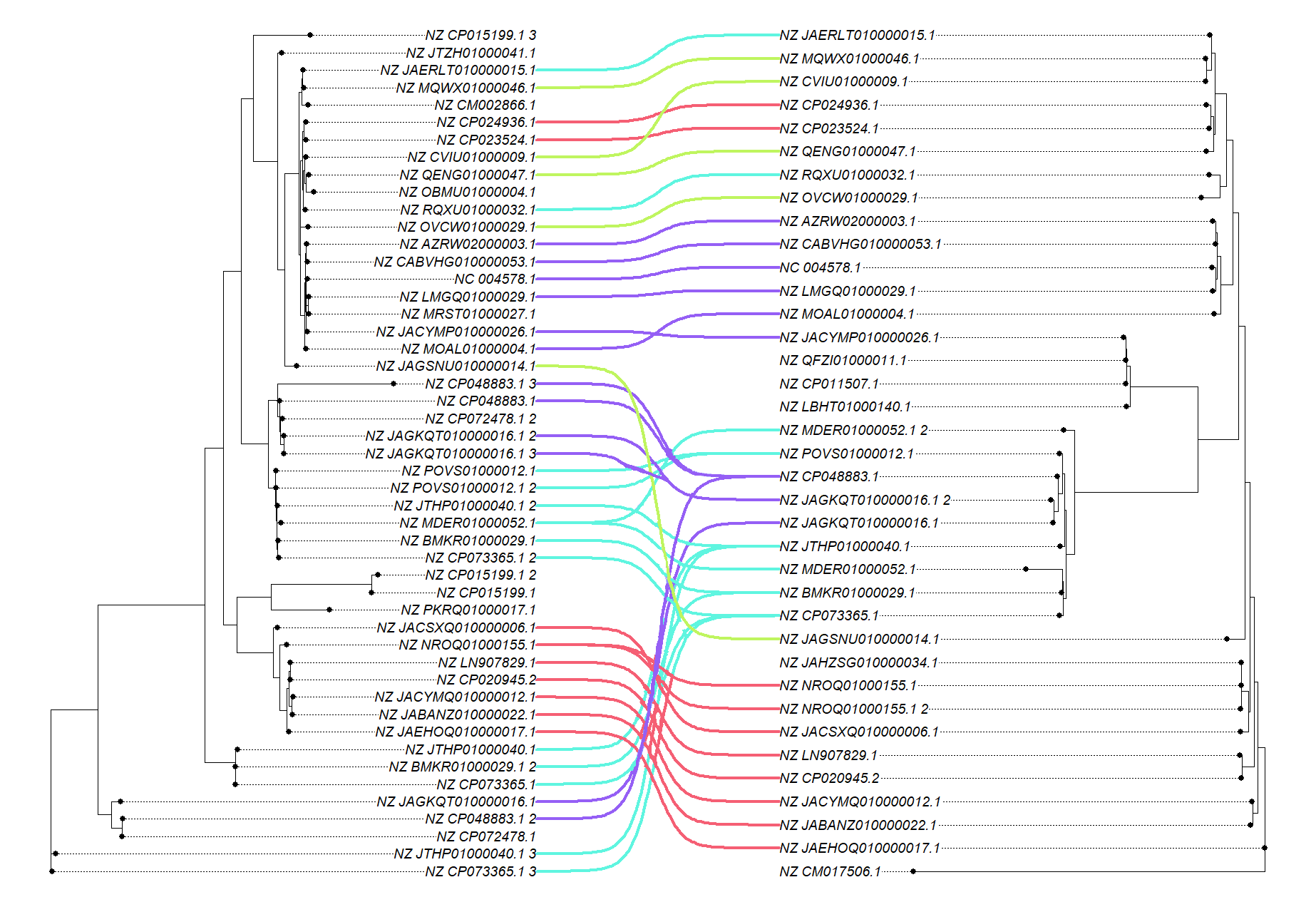

### altRvsaltA.png

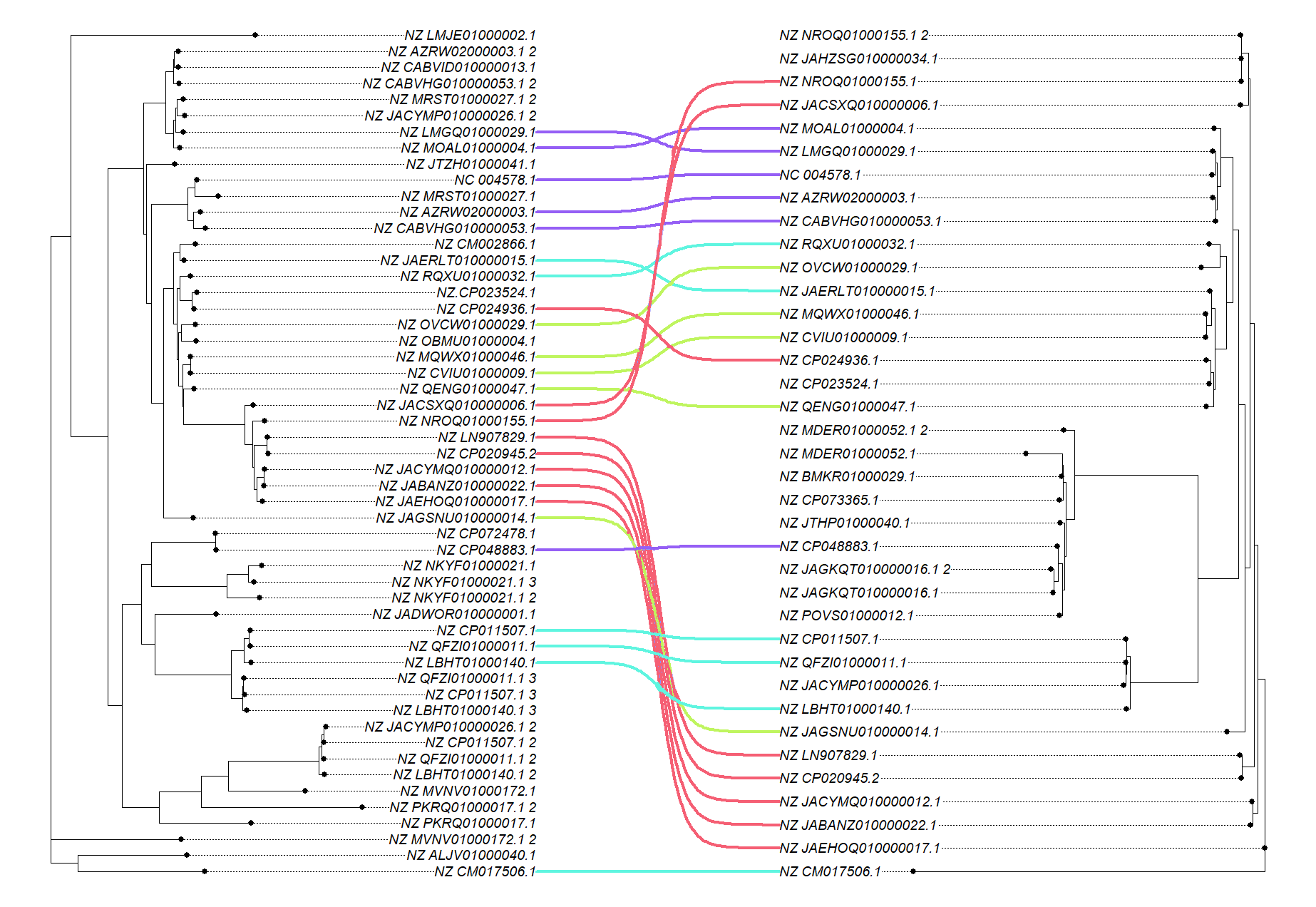

### altRvsaltE.png

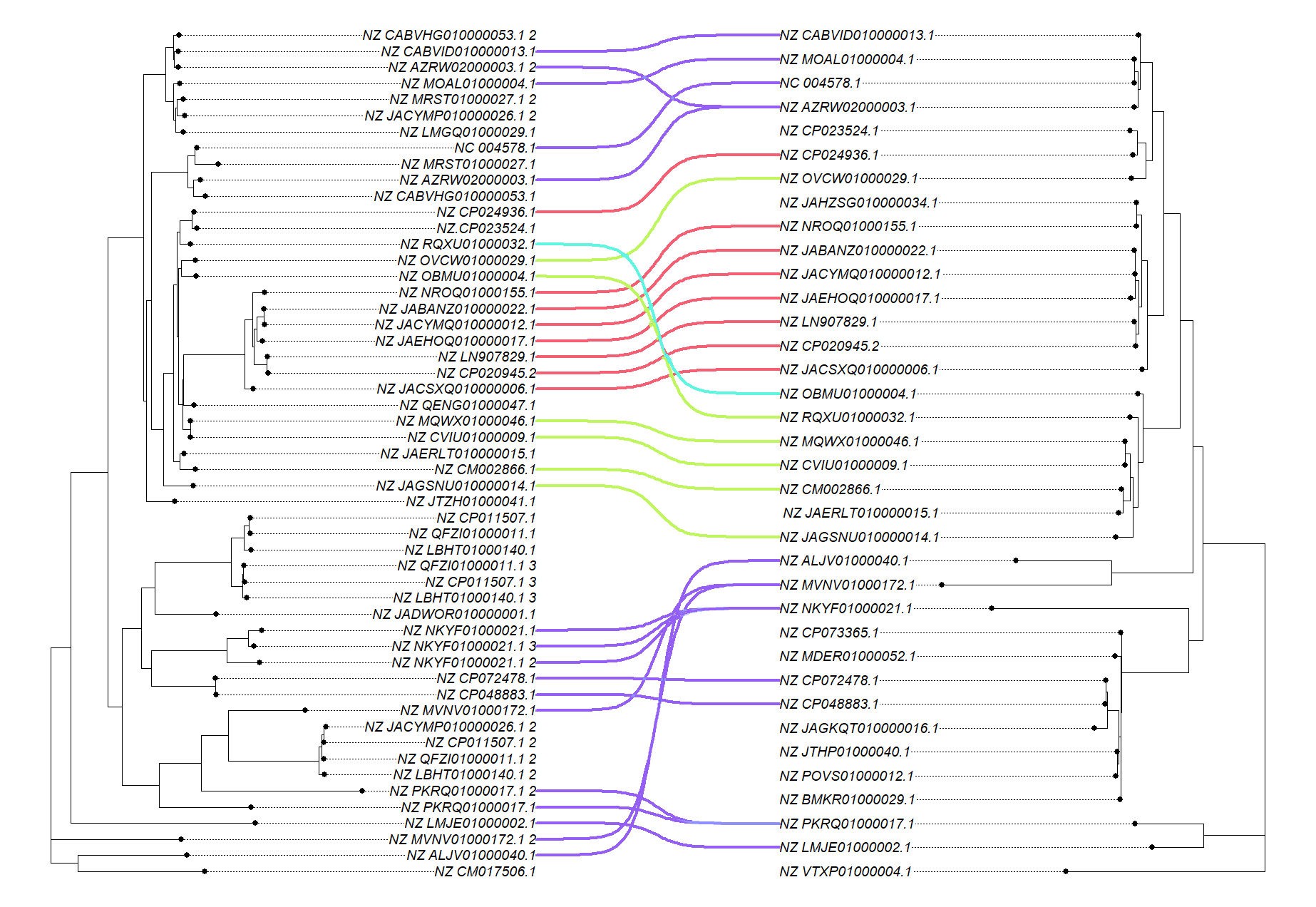

### altRvsaltJ.png

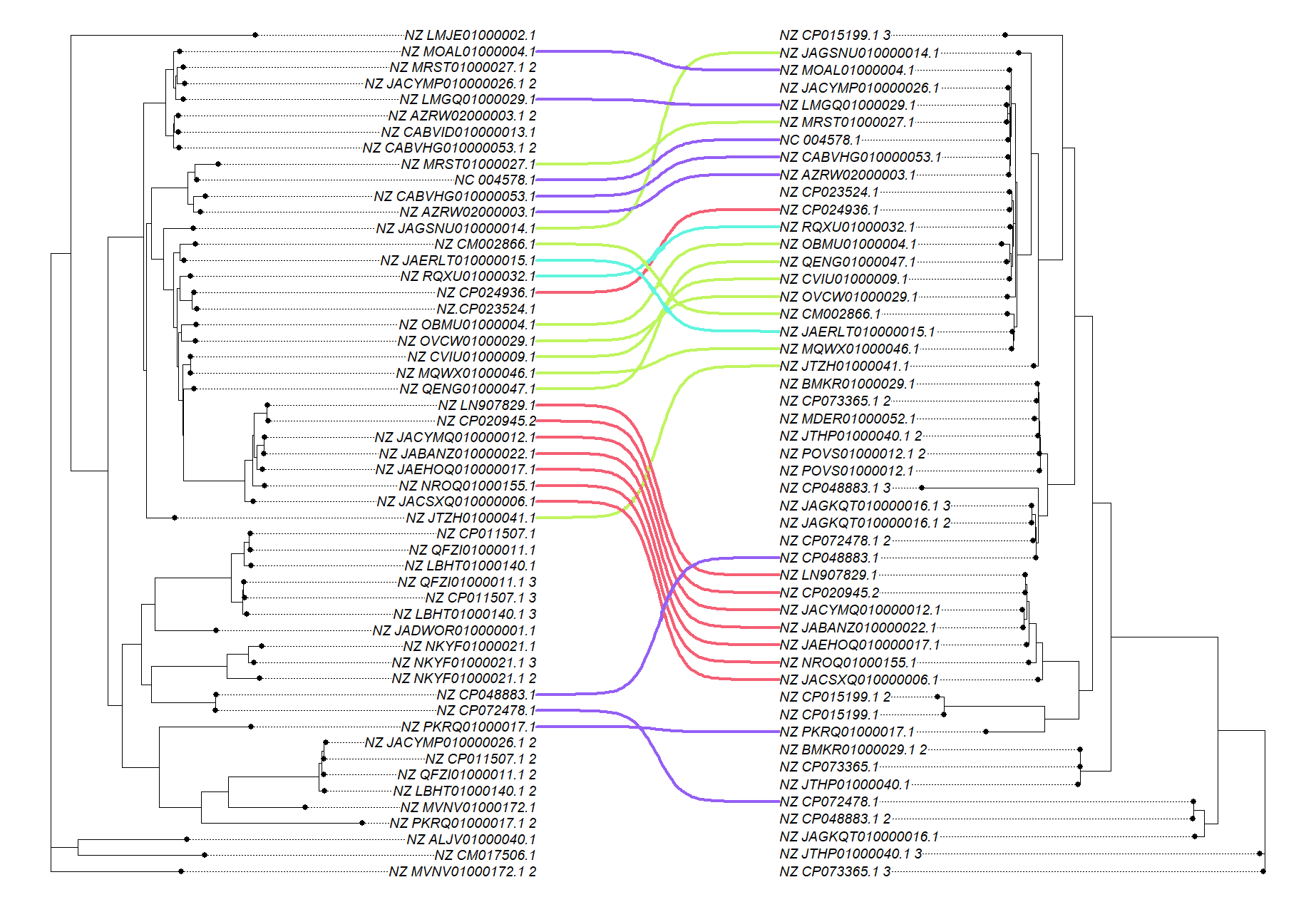

### Supplementary figure 3.png

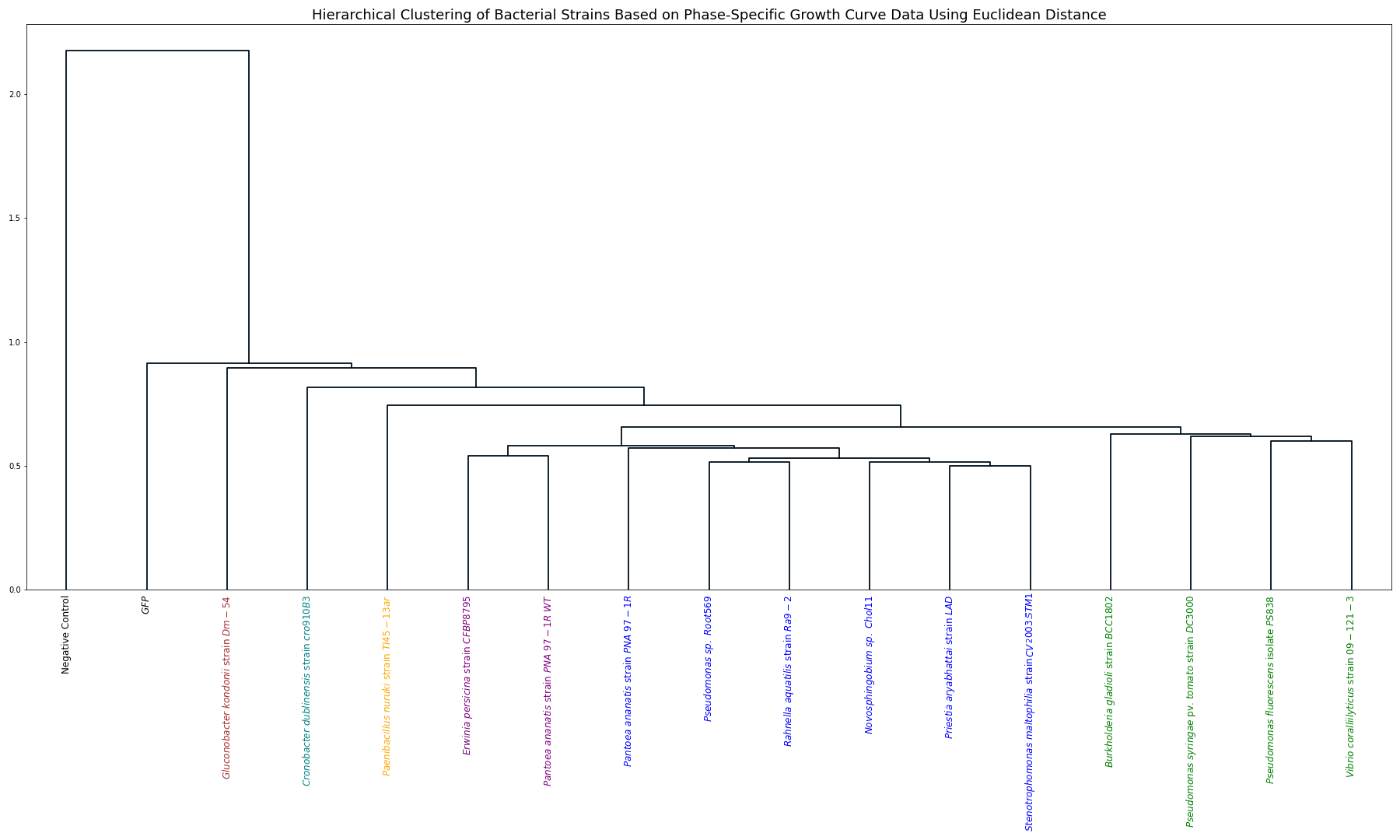
